# Identification of *Lactuca sativa* transcription factors impacting resistance to *Botrytis cinerea* through predictive network inference

**DOI:** 10.1101/2023.07.19.549542

**Authors:** Harry Pink, Adam Talbot, Ryan Carter, Richard Hickman, Oliver Cooper, Rebecca Law, Gillian Higgins, Chenyi Yao, Frances Gawthrop, Paul Hand, David Pink, John Clarkson, Katherine Denby

## Abstract

Lettuce is susceptible to a wide range of plant pathogens including the fungal pathogens *Botrytis cinerea* and *Sclerotinia sclerotiorum*, causal agents of grey mould and lettuce drop, respectively. Chemical control is routinely used but there is an urgent need to develop varieties with enhanced resistance given the economic and environmental costs of preventative pesticide sprays, the prevalence of fungicide-resistant isolates of both pathogens in the field, and the increasing withdrawal of approved fungicides through legislation. Resistance against *Botrytis cinerea* and *Sclerotinia sclerotiorum* is quantitative, governed by multiple small-medium impact loci, with plant responses involving large-scale transcriptional reprogramming. The elucidation of the gene regulatory networks (GRNs) mediating these responses will not only identify key transcriptional regulators but also interactions between regulators and show how the defence response is fine-tuned to a particular pathogen. We generated high-resolution (14 time points) time series expression data from lettuce leaves following mock-inoculation or inoculation with *B. cinerea*, capturing the dynamics of the transcriptional response to infection. Integrating this data with a time series dataset from *S. sclerotiorum* infection of lettuce identified a core set of 4362 genes similarly differentially expressed in response to both pathogens. Using the expression data for these core genes (with additional single time point data from 21 different lettuce accessions) we inferred a GRN underlying the lettuce defence response to these pathogens. Using the GRN, we have predicted and validated key regulators of lettuce immunity, identifying both positive (LsBOS1) and negative (LsNAC53) regulators of defence against *B. cinerea*, as well as downstream target genes. These data provide a high level of detail on defence-induced transcriptional change in a crop species and a GRN with the ability to predict transcription factors mediating disease resistance both in lettuce and other species.

## Introduction

*Botrytis cinerea* is a devastating plant pathogen able to infect over 200 plant species including numerous important crop species, costing over $10 billion per year in control attempts and crop losses. *B. cinerea* can infect pre or post-harvest, and isolates have been identified which show resistance to fungicides (Rupp et al., 2016). *B. cinerea* is a generalist necrotrophic pathogen, secreting a vast arsenal of phytotoxins and cell wall degrading enzymes to induce cell death in its host (Williamson et al., 2007). *Sclerotinia sclerotiorum* is another necrotrophic fungal pathogen which is closely related to *B. cinerea* and employs similar infection strategies (Amselem et al., 2011). *Lactuca sativa* (lettuce) is a nutritionally and economically important crop species with a global value of $US2.4 billion. Lettuce is highly susceptible to a number of plant pathogens including the fungal pathogens,*B. cinerea* and *S. sclerotiorum*.

*Arabidopsis thaliana-B. cinerea* is one of the most extensively studied pathosystems in plant pathology, providing a high-level understanding of the complex plant-pathogen interactions. Upon *B. cinerea* infection, microbe-associated molecular patterns (MAMPs) or damaged associated molecular patterns (DAMPs) are recognised by pathogen recognition receptors (PRRs), which trigger downstream signalling to activate a defence response. For example, fungal cell wall component chitin acts as a MAMP, which is recognised by CERK1 (Chitin Elicitor Receptor Kinase 1) (Miya et al., 2007). Chitin-activated CERK1 phosphorylates PLB27, which in turn initiates a MAP kinase cascade via MAPKKK5 (Shinya et al., 2014). This recognition pathway is required for defence against multiple necrotrophic fungi including*B. cinerea* and *Alternaria brassicicola* (Yamada et al., 2016; Liu et al., 2018). Such signalling cascades trigger large-scale transcriptional reprograming changes to combat pathogen infection. In response to *B. cinerea*, Arabidopsis undergoes massive transcriptional reprogramming, differentially expressing over 9000 genes (Windram et al., 2012).

Pathogen-induced transcriptional reprogramming is significantly impacted by plant hormone signalling networks, which themselves exhibit multiple levels of crosstalk. Two key defence hromones, jasmonic acid (JA) and salicylic acid (SA), induce differential expression of thousands of genes in Arabidopsis (Hickman et al., 2017, 2019; Zander et al., 2020). Generally, JA and ethylene (ET) promote defence against necrotrophic pathogens, and SA against biotrophic pathogens, However, exogenous application of SA increases*B. cinerea* resistance (Ferrari et al., 2003) indicating that effective defence requires complex interaction between hormone networks to fine tune gene expression in response to different pathogens based on their lifestyle. Transcription factors (TFs) are central in integrating multiple hormone signals and finetuning the defence in specific scenarios (Nomoto et al., 2021; Aerts et al., 2021; Caarls et al., 2017; Ndamukong et al., 2007; Zhang et al., 2014; Tsuda and Somssich, 2015). There are several major TF families which coordinate these defence responses such as ERFs (Ethylene Responsive Factors), WRKYs, MYBs, NACs (NAM-ATAF1-CUC2 family) and bHLHs (basic helix-loop-helix) (Tsuda and Somssich, 2015). Many TFs within these families, such as ERF1 (Berrocal-Lobo et al., 2002), ORA59 (ERF)(Pre et al., 2008), WRKY33 (Birkenbihl et al., 2012; Liu et al., 2015), WRKY70/WRKY54 (Li et al., 2017), BOS1 (MYB)(Mengiste et al., 2003), MYC2 (bHLH) (Lorenzo et al., 2004) and NAC019/NAC055 (Bu et al., 2008) have been identified as key regulators of the defence response to *B. cinerea*, either promoting resistance or susceptibility to the fungus. What is not clear is how these individual TFs operate within gene regulatory networks (GRNs) to shape the defence response against a particular pathogen, or whether orthologues of these TFs (and/or different TFs) are important in lettuce defence against *B. cinerea*.

With recent advances in high-throughput sequencing and the availability of whole-transcriptome expression data it is possible to take a systems biology approach to identify regulators and understand the complex GRNs in which these regulators interact. Inference of GRNs can be performed to predict key regulators, or “hub genes”, which regulate other genes within a network. In causal network inference, nodes (genes) are linked by directional edges and expression of the upstream gene is predicted to impact expression of the downstream gene. This contrasts with co-expression networks where nodes are linked by an edge if they share a similar expression profile across the input data sets, with no prediction of upstream regulation. For causal network inference, high-resolution temporal transcriptome data is critical. We previously constructed a GRN based on an Arabidopsis time series gene expression data set (*B. cinerea-* and mock-inoculated samples, 24 time points) which identified TGA3 as a network ‘hub’ positively regulating*B. cinerea* resistance (Windram et al., 2012). However, incorporating multiple complementary datasets into network inference is likely to increase the accuracy of the resulting network model. Here, we present a high-resolution *B. cinerea* infection (and mock) time-series from a lettuce cultivar (*L. sativa* cv. Saladin). We construct a GRN integrating our *B. cinerea-*infection time-series data with three additional transcriptomic datasets from lettuce infected with B*. cinerea* and *S. sclerotiorum* (Pink et al., 2022; Ransom et al., 2023). Together, these data provide an unprecedented level of detail about the transcriptional defence response for a crop species, identify a core set of genes that respond to *B. cinerea* and *S. sclerotiorum*, and generates a high-confidence GRN model underlying the lettuce defence response. This GRN accurately predicts LsBOS1 and LsNAC53 as key regulators of defence against *B. cinerea*, as well as validated downstream target genes of LsNAC53.

## Results

### Shared transcriptional reprogramming in lettuce during *Botrytis cinerea* and *Sclerotinia sclerotiorum* infection

We profiled the transcriptome of lettuce leaves (cv. Saladin) following inoculation with spores of *Botrytis cinerea* pepper isolate (Windram et al., 2012). The third leaf from 4-week old plants was inoculated with four droplets of 5 x 10^5^ mL spore suspension or mock control. One cm diameter leaf discs around each inoculation site were harvested every 3 hours between 9 and 48 hours post-inoculation (hpi) with the four discs from one leaf pooled as one sample (Figure 1). Three leaves were sampled at each of the 14 time points, for both inoculated and mock, as biological replicates. Total RNA was extracted from each sample and mRNA profiled using Illumina short read sequencing and reads mapped to a combined lettuce-*B. cinerea* transcriptome (Supplementary Dataset 1). As expected, the proportion of reads mapping to the *B. cinerea* genome increased over time (Figure 2a) although even at the later time points the vast majority of reads were mapping to lettuce transcripts (Supp Dataset 1). After quantification of reads (Supp Dataset 2), principal component analysis of lettuce gene expression highlighted the similarity between biological replicates and showed clear differences in lettuce gene expression as infection progressed (Supp Fig 1). As expected, we observe diel oscillation in expression of some genes across the time series demonstrating the need for a mock-inoculated time series as opposed to a single time-point control (Supp Fig 2). Differential expression analysis was performed using limma-voom (Ritchie et al., 2015) revealing 6713 differentially expressed genes (DEGs) over the time series, 3524 up- and 3189 down-regulated (Supp Dataset 3, Supp Figure 3). Hence, in the hours leading up to and during initial visible lesion development, there is large-scale transcriptional reprogramming in lettuce leaves in response to *B. cinerea* inoculation.

**Figure 1:**
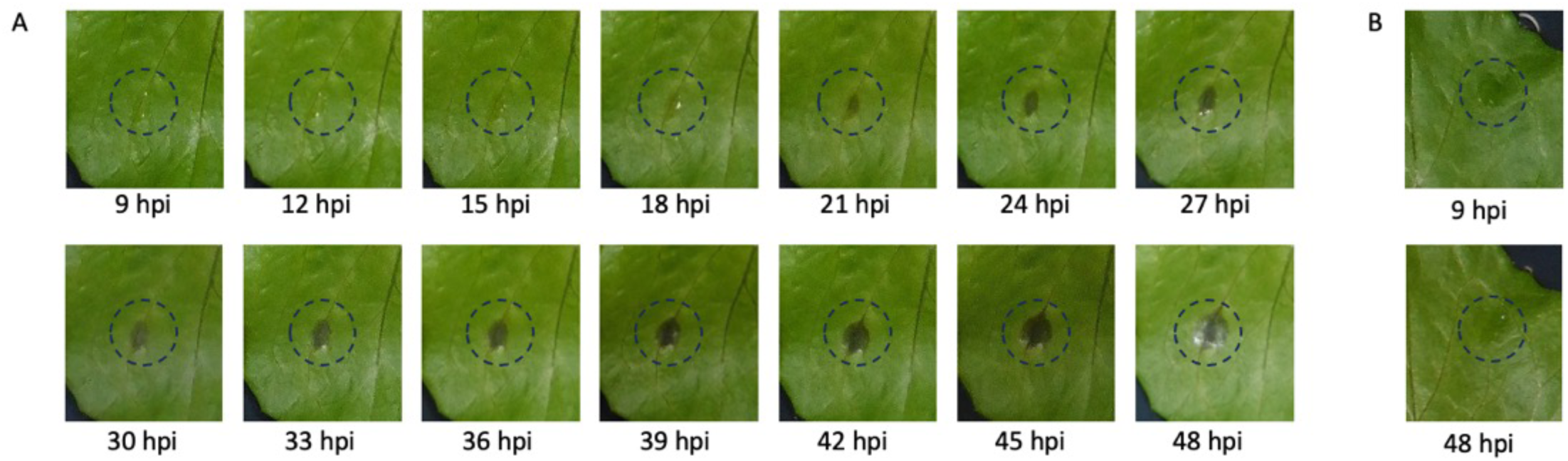
Time series of *Botrytis cinerea* infection on lettuce leaves. A) 10 μL droplets of a suspension of *B. cinerea* spores (5 x 10^4^ spores mL^-1^) were placed on detached leaves from 4 week old lettuce cv. Saladin plants. Images show the same leaf every 3 hr from 9 hours post inoculation (hpi) to 48 hpi. B) A mock inoculated leaf at 9 hpi and 48 hpi. The dashed line indicates the size of the 1 cm diameter disc used for sampling.

**Figure 2:**
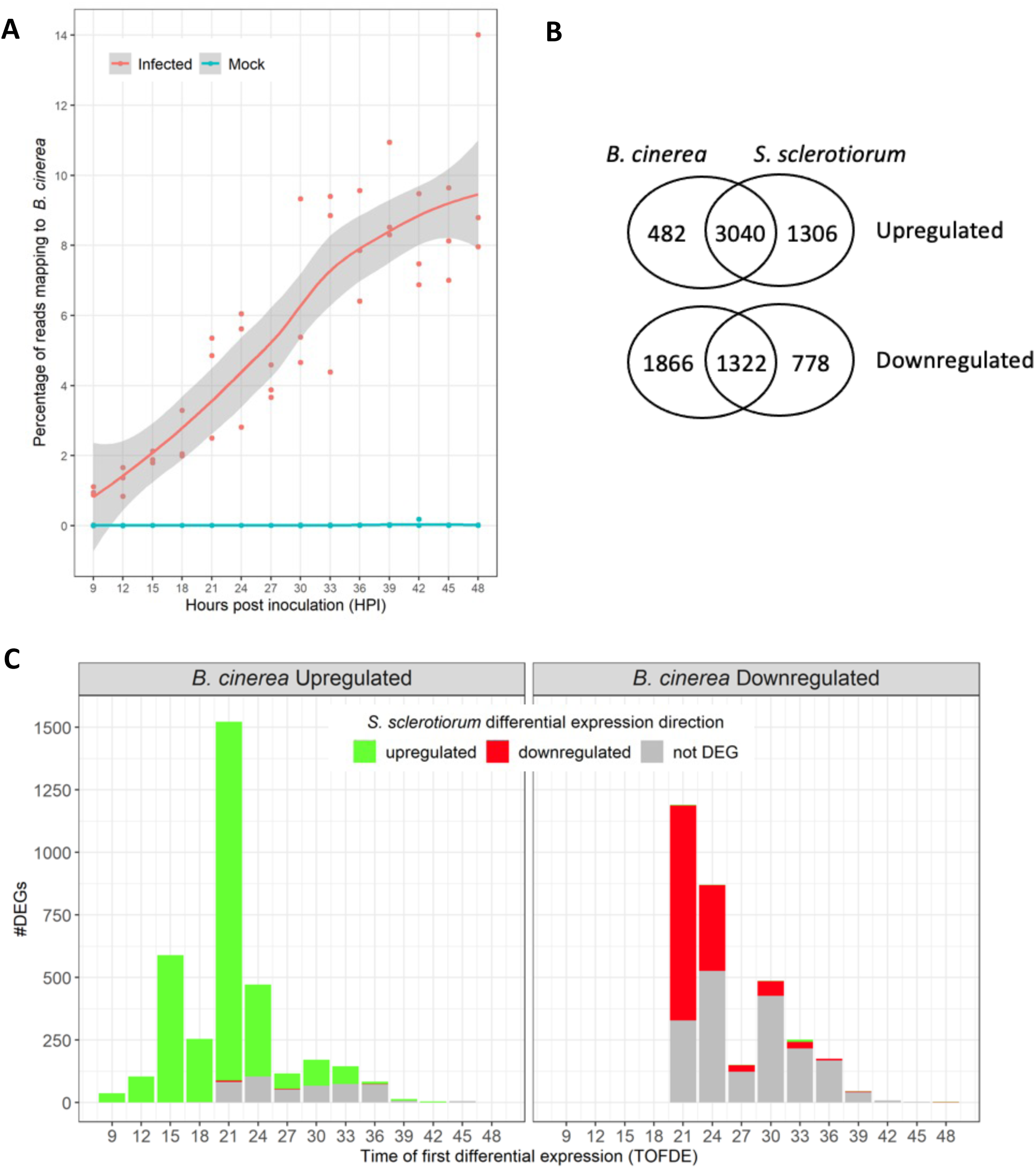
*B. cinerea* infection leads to large-scale transcriptional reprogramming in lettuce. (A) The percentage of RNAseq reads mapping to the *B. cinerea* transcriptome compared to the total number of mapped reads in each times series sample. Individual data points are shown along with a smoothed regression line and 95% confidence interval, in grey. As expected, there are no or extremely low numbers of reads mapping to *B. cinerea* in the mock-inoculated samples. The proportion of reads in each sample mapping to the *B. cinerea* transcriptome increased over time after inoculation, an indication of pathogen growth during the infection. Red indicates infected samples and blue, mock. (B) Up- and down-regulated lettuce genes following inoculation with *B. cinerea (this study)* and *S. sclerotiorum* (Ransom et al 2023). 4362 genes are differentially expressed after inoculation with both pathogens with the same direction of expression change. (C) Timing of First Differential expression (TOFDE) of lettuce DEGs during *B. cinerea* infection, separated into upregulated (left panel) or downregulated (right) genes. The number of DEGs with a TOFDE at each time point is indicated. Colouring indicates whether the DEGs are upregulated, downregulated or not differentially expressed (not DEG) in response to *S. sclerotiorum* infection (Ransom et al. 2023).

Previously, we conducted an independent RNAseq time-series, capturing the transcriptome response in lettuce to inoculation with *S. sclerotiorum*, a fungal pathogen closely related to *B. cinerea* (Ransom et al., 2023). This analysis identified a similar number of DEGs compared to mock-inoculated controls (6446; 4346 up-regulated, 2100 down-regulated). There is a striking similarity between the transcriptional reprogramming occurring in lettuce in response to *B. cinerea* and *S. sclerotiorum*, with 4390 DEGs common to both. Furthermore, the changes in gene expression are overwhelmingly in the same direction (3040 DEGs upregulated and 1322 DEGs downregulated in response to both *B. cinerea* and *S. sclerotiorum*, with only 28 genes showing an opposite change in expression). This reveals a core set of 4362 DEGs which have the same direction of differential expression in response to both pathogens (Figure 2B, Supp Dataset 4).

To determine critical periods for transcriptional reprogramming in response to these generalist necrotrophic pathogens, we identified the time of first differential expression (TOFDE) for all lettuce DEGs during *B. cinerea* infection (Supp Dataset 4). The early phase of transcriptional reprogramming (9-18 hpi) is dominated by upregulation of DEGs (Fig 2C) with 99.6% (982/986) of these early DEGs are also upregulated in response to*S. sclerotiorum,* indicating an early conserved defence response. The vast proportion of DEGs after*B. cinerea* inoculation are first differentially expressed at 21 and 24 hpi (60%: 4055/6713). The late phase of transcriptional reprogramming (27-48 hpi) consists of more down-regulated DEGs than upregulated and has the least overlap with genes differentially expressed during *S. sclerotiorum* infection (Figure 2C). At least part of the reduced overlap for DEGs with later TOFDE is likely due to the slower progress of infection by *S. sclerotiorum* compared to *B. cinerea*, with the lettuce-*B. cinerea* downregulated genes typically having later TOFDE and the lettuce-*S. sclerotiorum* time series capturing less of the later response.

### Conserved and species-specific defence responses in lettuce and Arabidopsis

The Arabidopsis-*B. cinerea* pathosystem has been extensively characterised over the last 20 years and is now well-understood. Although lettuce (Asterid) and Arabidopsis (Rosid) are distant species (diverging approx. 125 million years ago)(Zeng et al., 2017), we expect there to be similarities in defence strategies against both *B. cinerea* and *S. sclerotiorum*, given the broad host range of these pathogens. Time-series transcriptome profiling of the Arabidopsis defence response against *B. cinerea* was previously carried out using microarrays capturing gene expression from 2 to 48 hpi (Windram et al., 2012). There is overlap between the DEGs in lettuce and Arabidopsis after *B. cinerea* inoculation (Supp Figure 4) with approx. 30% and 43% respectively of up- and down-regulated lettuce DEGs with their Arabidopsis orthologue differentially expressed in the same direction. However, a significant proportion of DEGs in lettuce are orthologous to Arabidopsis DEGs with an opposite direction of expression change, or which do not change in expression during *B. cinerea* infection of Arabidopsis. This suggests that there are both conserved and species-specific aspects to the defence response.

**Figure 3:**
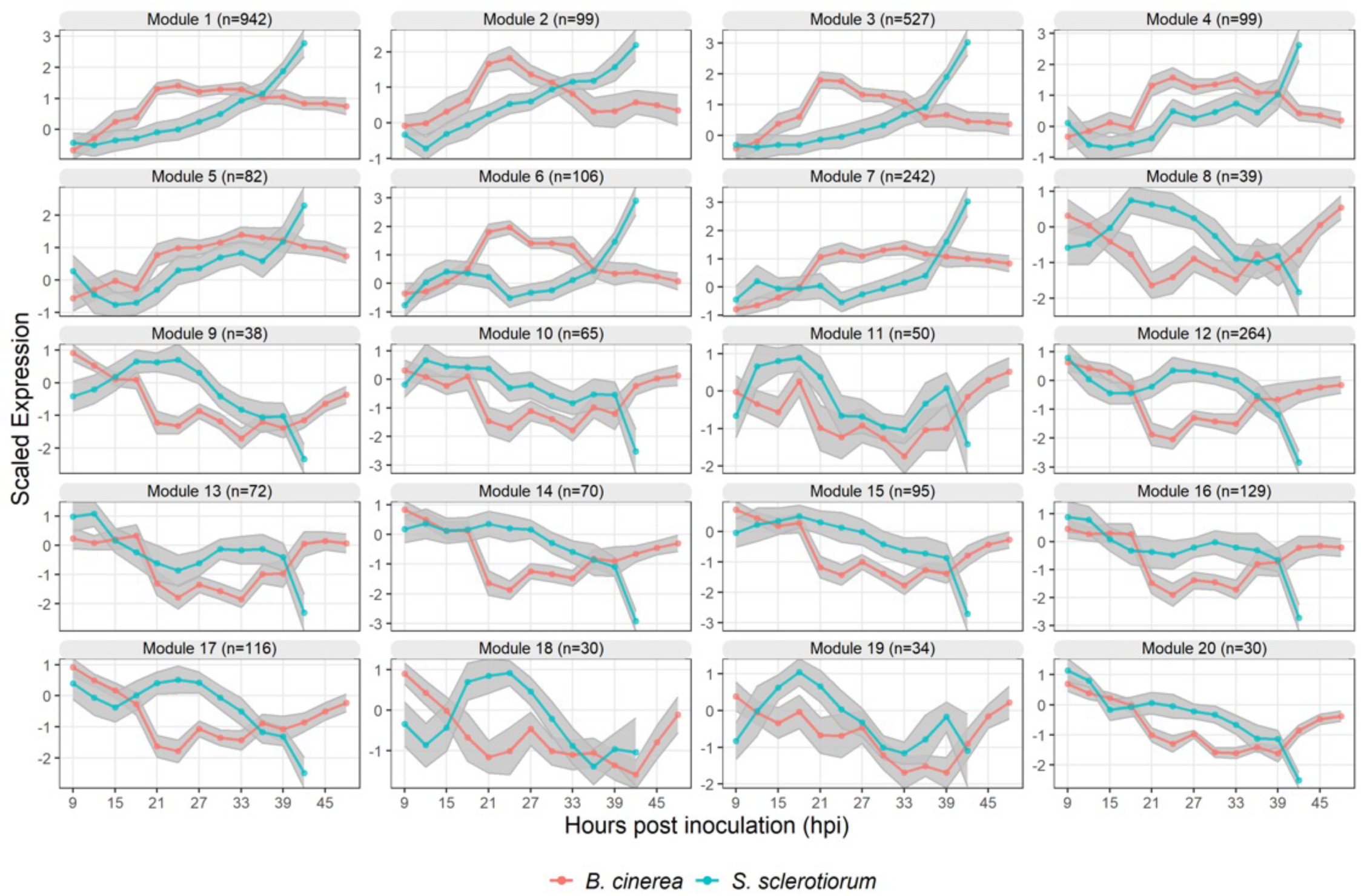
Modules of lettuce DEGs co-expressed following both *B. cinerea* (red) and *S. sclerotiorum* (blue) infection. 3129 of the 4362 lettuce genes differentially expressed in both the *lettuce-B. cinerea* and lettuce-*S. sclerotiorum* time-series are included in a module, with the mean scaled log2 expression profile (solid line) and 95% confidence interval (grey area) of the genes in each module shown. N represents the number of genes within a module. Modules 1 – 7 contain upregulated DEGs, and modules 8 to 20 contain downregulated DEGs (compared to mock–inoculated control in each time series).

**Figure 4:**
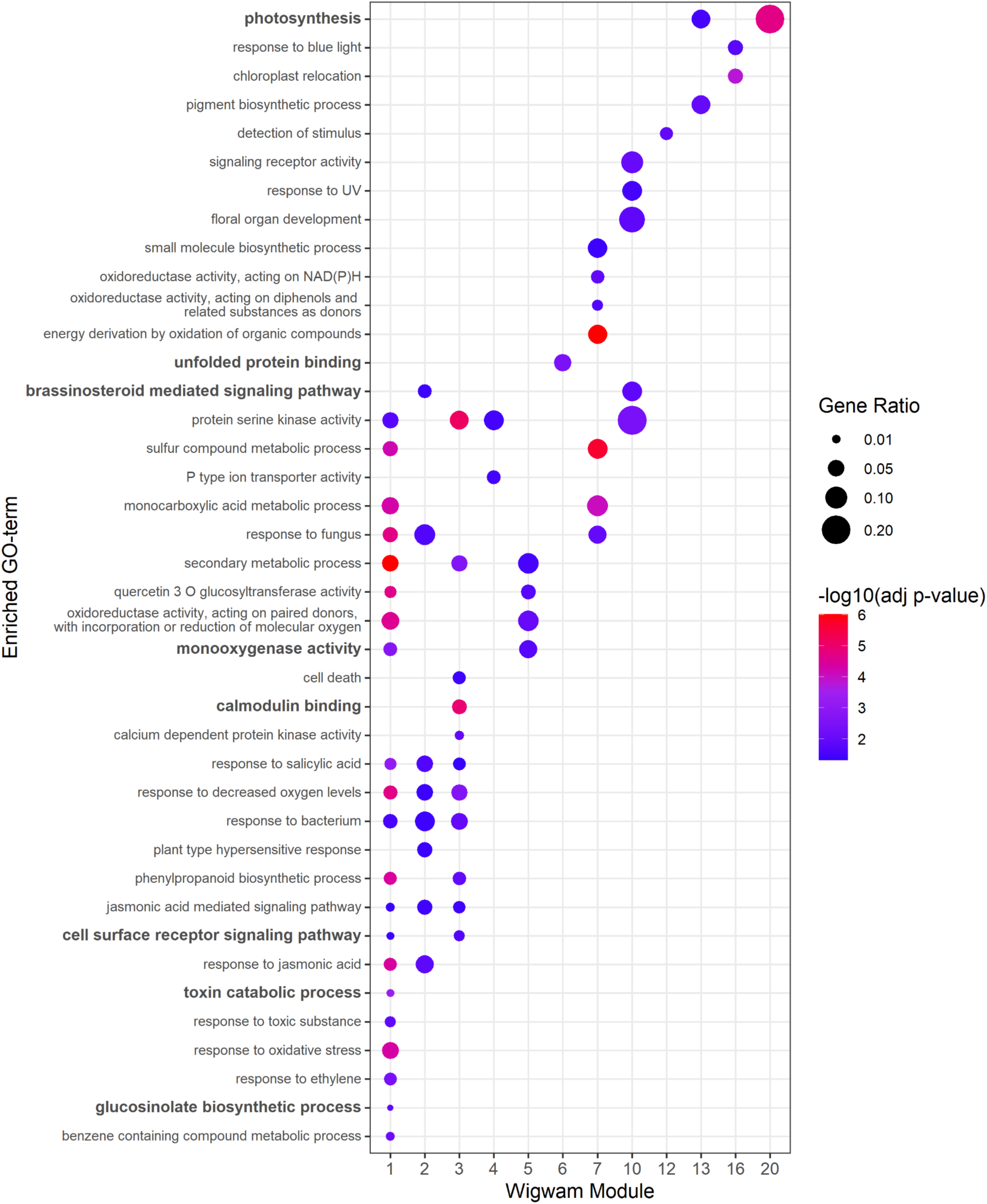
Modules of co-expressed lettuce genes are enriched for genes involved in different biological processes. Selected gene ontology (GO) terms significantly enriched in modules of lettuce genes co-expressed in response to *B. cinerea* and *S. sclerotiorum* are indicated, with colour indicating the statistical significance of the enrichment, and the size of the circle indicating the scale of enrichment for that term. Enrichment analysis was carried out using annotations of the Arabidopsis orthologue of each lettuce gene (where available) against a background of Arabidopsis orthologues of all lettuce genes detected in the mock- *or B.* cinerea-inoculated time series.

Many lettuce orthologues of well characterised Arabidopsis regulators with a known role in defence against *B. cinerea* were identified as DEGs in both time-series data sets (Supp Fig 5), including genes involved in JA and ET signalling (*LsERF1*, *LsMYC2*, *LsWRKY33*), JA and ET biosynthesis (Allene oxide synthase, *LsAOS*; lipoxygenase 1, *LsLOX1*), SA signalling (*LsWRKY54*, *LsWRKY70*, Enhanced disease susceptibility 1, *LsEDS1*) and SA biosynthesis (*LsICS2*). WRKY54, a known SA regulator, has two lettuce orthologues differentially expressed, one of which is upregulated, the other downregulated. The expression profiles of these genes also illustrate the slower progression of infection by *S. sclerotiorum* with changes in gene expression delayed compared to the *B. cinerea* time series.

**Figure 5:**
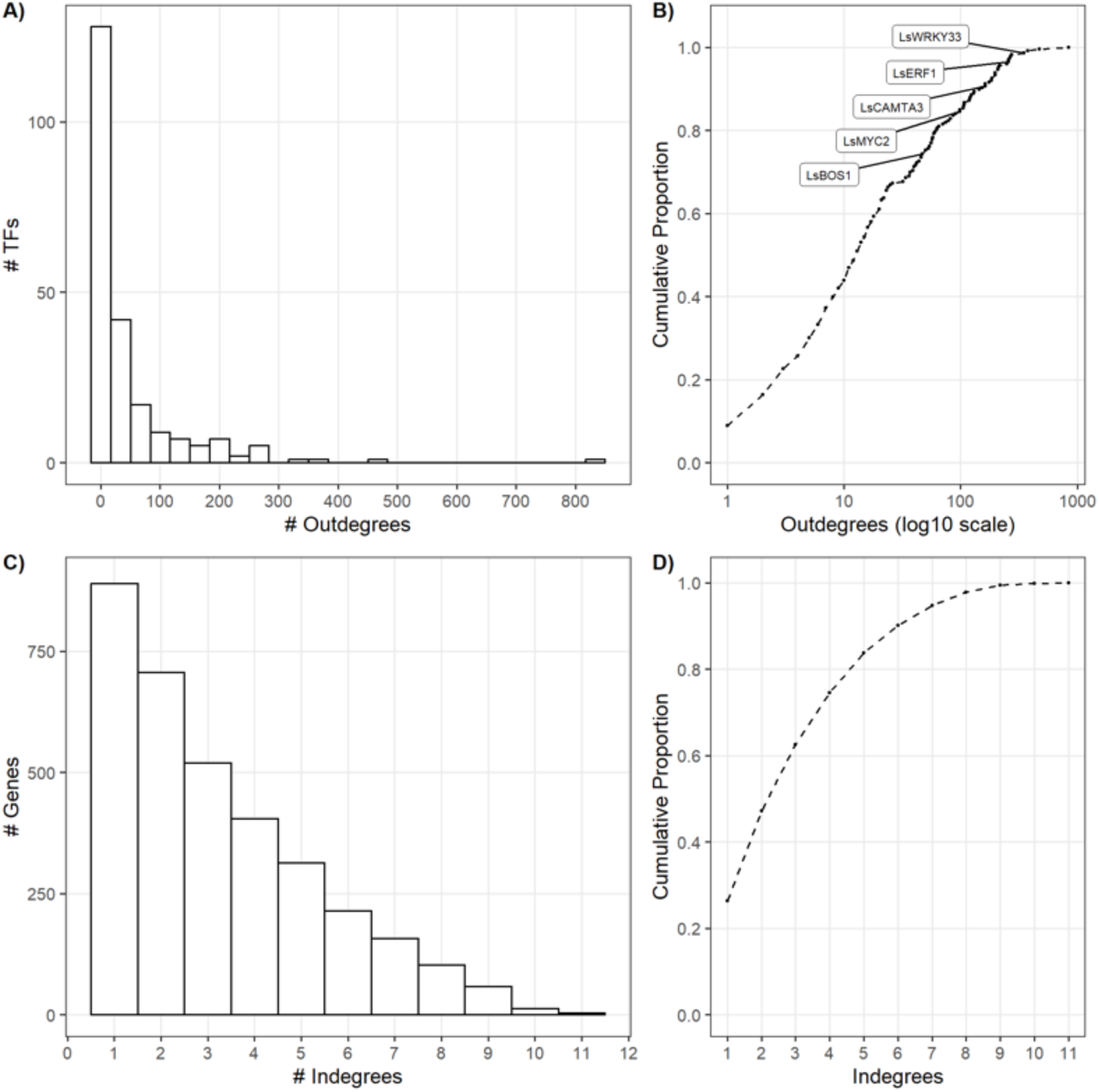
Characteristics of the gene regulatory network predicted to mediate the lettuce transcriptional response to *B. cinerea* and *S. sclerotiorum* infection. A) The distribution of transcription factor outdegree (i.e. number of downstream target genes); B) Empirical Cumulative Distribution Function (ECDF) of transcription factors outdegree, showing the proportion of transcription factors with >X outdegrees. Known regulators mentioned in the text are highlighted; C) The distribution of indegrees for each target gene in the network (i.e. the number of transcription factors predicted to regulate a gene); D) ECDF of node indegree, showing the proportion of genes in the network with >X indegrees.

A striking species difference is expression of plant defensins (PDFs), some of which are key marker genes of the JA-activated defence pathway in Arabidopsis (Brown et al., 2003; Manners et al., 1998). *AtPDF1.2* shows dramatic upregulation in response to *B. cinerea* and *Alternaria brassicicola* infection, which is abolished in both *coi1-1* mutants and ORA59 RNAi lines (Pre et al., 2008). *AtPDF1.1* and *AtPDF1.3* also show upregulation in response to *B. cinerea* (Ingle et al., 2015). We identified 13 putative PDFs in lettuce (LsPDFs), which contain the gamma-thionin domain (Pfam PF00304) and were shorter than 150 amino acids in length. Phylogenetic analysis of these with Arabidopsis defensins (AtPDFs), and characterised plant defensins with anti-fungal activity from other species (Lacerda et al., 2014)(Supp Figure 6a) indicated the similarity of the putative lettuce defensins to these proteins, particularly to AtPDF families 1 and 2. However, only 8 *LsPDF* genes had detectable expression in leaves and none were upregulated after *B. cinerea* infection (Supp Fig 6b). Five predicted LsPDFs have very low levels of expression in our samples, two show constitutively high levels of expression, and Lsat_1_v5_gn_5_70941 is downregulated after*B. cinerea* infection. It is possible that there is sufficient anti-fungal activity from the defensin genes with high levels of expression or that the pathogen may be preventing the upregulation of *LsPDFs*, or potentially driving reduction of Lsat_1_v5_gn_5_70941 mRNA through effector molecules introduced into the plant.

**Figure 6:**
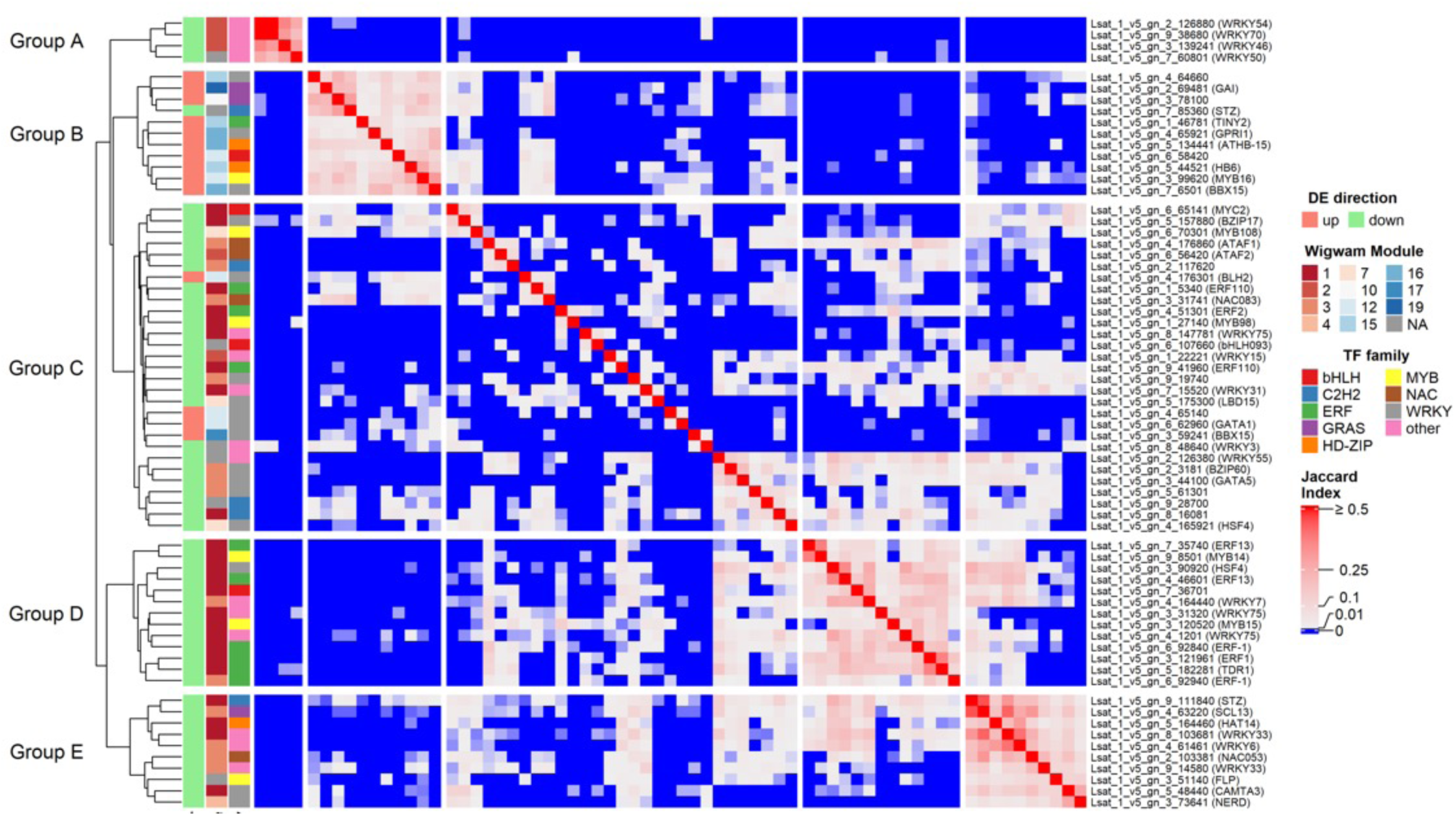
Pairwise similarity of predicted target genes of lettuce GRN TF hubs. The **h**eatmap shows the pairwise Jaccard Index (proportion of overlap) of the predicted targets of all lettuce hub TFs (≥40 predicted targets). Rows and columns are clustered on Euclidian distance. Row and column annotations indicate the TF family and time-series module of each hub gene, as well as the direction of differential expression of the hub gene following infection by *B. cinerea* or *S. sclerotiorum*.

### Co-expression modules highlight specific biological functions and processes during infection

We used the Wigwams (Polanski et al., 2014) algorithm to identify non-redundant modules of lettuce genes co-expressed in both the *B. cinerea* and *S. sclerotiorum* infection time-series. Wigwams does not force every gene into a module, unlike typical clustering algorithms, and evaluates each putative module for statistical significance in an attempt to identify co-expression due to true co-regulation, rather than simply because of the frequency of a particular expression profile. 3129 (72%) of the common 4362 DEGs were grouped into 20 co-expressed modules, with a median size of 90 genes (Figure 3, Supplementary dataset 5a). Two of the modules are very large (module 1 and 3), containing 942 and 527 genes respectively.

It is clear from the expression profiles of these modules, particularly modules 1 to 7, that the lettuce response to *B. cinerea* infection is faster than that to *S. sclerotiorum*, with a significant transcriptional shift by 21 hpi for *B.* cinerea and only happening by the end of the time series (42 hpi) for *S. sclerotiorum.* In the *B. cinerea* time series, several modules show transient upregulation of mRNA levels (e.g. modules 2 and 6) while others (e.g. modules 1 and 7) have increased expression levels that last throughout the time series. However, expression of all the downregulated modules during *B. cinerea* infection starts to recover either immediately after 21 hpi (e.g. modules 12 and 14) or from 39 hpi (e.g. modules 9, 15 and 19). Many of these modules show expression profiles still decreasing in the*S. sclerotiorum* time series.

We tested whether these modules were enriched for specific biological functions using gene ontology (GO) enrichment performed using annotations of the Arabidopsis orthologues of lettuce genes within each module. Arabidopsis orthologues of all the lettuce genes detected in the mock or *B. cinerea*-inoculated time series were used as the background set. 13 modules were significantly enriched for genes with a particular GO term (Supp Table 5b). Additionally, we performed protein domain enrichment, using Pfam and Panther annotations, again comparing against a background set of all genes with detectable expression in either time-series (Supp Dataset 5c).

General defence-related GO terms were enriched across multiple modules, with “response to fungus”, “secondary metabolic processes” and “response to bacterium” being significantly over-represented in 3 modules each (Figure 4). This was expected, as general responses like these are unlikely to be limited to a single group of genes. However, we also identified GO terms enriched in a single module, suggesting that these modules contain distinct biologically-relevant groups of genes. Module-specific functions include response to ethylene (module 1), cell death (module 3), unfolded protein binding (module 6), response to UV (module 10), pigment biosynthetic process (module 13) and chloroplast relocation (module 16).

Module 1 is a large group of 942 genes, accounting for 21.6% of all common*B. cinerea* /*S. sclerotiorum* DEGs and 30.0% of all DEGs assigned to a module, demonstrating that despite the integration of two high-density time series datasets, large proportions of the transcriptome appear to change within a very short time-frame. Enrichment for biological functions associated with JA and ET signalling (known regulators of defence against*B. cinerea* in Arabidopsis) was evident. The GO terms “response to jasmonic acid”, “jasmonic acid biosynthetic process”, “response to ethylene” and “ethylene-activated signalling pathway” as well as AP2/ERF protein domains all show their highest levels of overrepresentation in module 1 (Figure 4, Supp dataset 5b). Lettuce orthologues of many key Arabidopsis JA/ET biosynthetic and/or response genes are identified in this module: 17 ERF domain TFs (including ERF1, three ERF13 orthologues and three ERF-1 orthologues), WRKY33, MYC2, LOX2 and EFE (ethylene forming enzyme). Downstream genes responding to JA/ET signalling have not yet been characterised in lettuce, however the large group of genes that are co-expressed with known JA/ET regulators and biosynthetic enzymes in module 1 may represent such genes in lettuce. The biosynthesis of key Arabidopsis phytoalexins such as camalexin and glucosinolates is JA/ET regulated (Hickman et al., 2017; Zhou et al., 2022) and promotes resistance to *B. cinerea* (Ferrari et al., 2003; Denby et al., 2004; Kliebenstein et al., 2005) and *S. sclerotiorum* (Stotz et al., 2011; Zhang et al., 2015).

Module 1 shows enrichment of cytochrome P450 protein domains as well as “glucosinolate biosynthetic process”, “isoprenoid metabolic process” and “secondary metabolic process” GO terms. Module 1 also contains three orthologues of PDR12, a transporter responsible for secreting camalexin in Arabidopsis (He et al., 2019). While camalexin is a phytoalexin specific to Arabidopsis, these transporters may be secreting other anti-fungal compounds in lettuce.

Lettuce is known to synthesise a diverse range of sesquiterpene lactones (STLs), including sulfate, oxalate and amino acid conjugates (Yang et al., 2022) with one compound, lettucenin A, shown to have anti-fungal activity against *B. cinerea* in vitro (Bennett et al., 1994; Sessa et al., 2000a). Germacrene A synthase (Bennett et al., 2002; Kwon et al., 2022), germacrene A oxidase (Nguyen et al., 2010) and costunolide synthase (Ikezawa et al., 2011) catalyse the production of costunolide, a key precursor of STLs. Genes encoding all three of these enzymes are in module 1 along with 37 additional cytochrome P450 encoding genes. Downstream of costunolide there is significant diversity in STL structures, with the biosynthetic pathways unknown. Hence these uncharacterised P450s, co-expressed with known STL biosynthetic genes, are good candidates for roles in the synthesis of STLs in lettuce. The presence of STL biosynthetic genes in module 1, may further suggest STL biosynthesis in lettuce is JA/ET regulated.

In addition to production of anti-microbial compounds, defence against *B. cinerea* is likely to require detoxification of pathogen toxins, including botrydial and botcinic acid (Zhang et al., 2019). The GO terms “Toxin catabolic process” along with Glutathione S-transferase (GST) and Aflatoxin B1 aldehyde reductase (AFAR) protein domains are overrepresented in module 1 genes. Aflatoxin B1 a mycotoxin produced by the saprophytic fungus *Aspergillus flavus* during infection of maize and peanut, is detoxified by AFAR enzymes (Judah et al., 1993; Klich, 2007). Although *B. cinerea* and *S. sclerotiorum* do not produce aflatoxin, the lettuce AFAR-like enzymes may have roles in detoxification of other pathogen-derived metabolites, with their presence in module 1 suggesting potential JA/ET regulation.

Module 3 (527 genes) is enriched for genes involved in pathogen perception, annotated with GO term “cell surface receptor signaling pathway”, as well as being the only module enriched for “cell death”. Orthologues of chitin-binding pathogen recognition receptors (PRRs) are present in module 3 such as lettuce orthologues of chitin-elicited receptor kinase 1 (CERK1) and lysin motif (LysM) receptor kinase 4 (LYK4), both of which have been shown to play a role in resistance to *B. cinerea* in Arabidopsis (Liu et al., 2018; Cao et al., 2014). A lettuce orthologue of SOBIR1 is also present in this module, along with orthologues of a receptor-like protein (RLP) and SERK4/BKK1 (BAK1-LIKE 1). SOBIR1 is a PRR known to promote *B. cinerea* and *S. sclerotiorum* resistance via recognition of elicitor peptides in co-receptor complexes with BAK1 and RLPs (Zhang et al., 2013; Albert et al., 2015, 2019; Ono et al., 2020), with SERK4/BKK1 (BAK1-LIKE 1) having functional redundancy with BAK1 (He et al., 2007; Schoonbeek et al., 2022). The presence of lettuce orthologues for these known pathogen recognition complexes in a single module suggests co-regulation of these genes in response to initial pathogen perception. Interestingly, a lettuce orthologue of BIR1 is also present in module 3. BIR1 negatively regulates the SOBIR1-BAK1 interaction (Liu et al., 2016; Ma et al., 2017) indicating coordinated regulation of mechanisms to dampen plant defence responses, potentially balancing effective defence with the physiological impact on the host plant. In the same co-expression module, we see overrepresentation of EF-hand protein domain and the presence of 5 genes encoding calcium-dependent protein kinases (CPKs), suggesting an important role for calcium signalling. Arabidopsis CPK mutants, *cpk1-1* and *cpk5/6/11*, show hyper-susceptibility to *B. cinerea*, with *cpk5/6/11* also showing a reduced response to oligogalacturonide DAMPs (Coca and Segundo, 2010; Gravino et al., 2015).

Module 6 is enriched for the GO term “unfolded protein response” (UPR), a response triggered by the accumulation of misfolded proteins in the endoplasmic reticulum, inducing chaperone expression to maintain correct protein folding (Bao and Howell, 2017). UPR has been shown to promote resistance to *Alternaria alternata*, a necrotrophic fungus, in *Nicotiana attenuata* (Xu et al., 2019). Lettuce orthologues of key chaperone proteins including ERdj3B, CNX1 and HSP89.1 (Liu et al., 2017; Song et al., 2010) are present in module 6.

Amongst the downregulated clusters, multiple photosynthesis and growth-related GO terms are significantly enriched, indicating a switch from growth to defence during necrotrophic pathogen infection. This has been seen in many plant defence responses to pathogens including during *B. cinerea* infection of Arabidopsis (Windram et al., 2012) and includes genes involved in chlorophyll synthesis (module 13), chloroplast localization (module 16) and photosynthesis reactions (module 20). Module 10 was significantly enriched for the GO term “brassinosteroid (BR) mediated signaling pathway”, containing orthologues of the BR receptor, BR insensitive 1 (BRI1) and a downstream signalling kinase, BR signalling kinase 2 (BSK2)(Tang et al., 2008). BRI1 is also a co-receptor of BAK1, and BR signalling inhibits BAK1-mediated immune signalling (Albrecht et al., 2012; Belkhadir et al., 2012), however, several BSKs have now been shown to interact with PRRs and promote *B. cinerea* resistance (Majhi et al., 2019, 2021).

### Conserved transcription factor DNA-binding motifs in gene modules

We would expect genes within a module with statistically significant co-expression to be co-regulated. We therefore tested for enrichment of known DNA-binding motifs in the promoters of the lettuce DEGs in each module. DNA sequence 1 Kb upstream from the transcriptional start site of all module genes was used as the putative promoter regions, and Arabidopsis DNA affinity purification sequencing (DAP-seq) data (O’Malley et al., 2016) used as the set of DNA binding motifs. Although this dataset characterises DNA binding motifs in Arabidopsis, not lettuce, we expect DNA-binding motifs to be conserved across species, and there is limited research on lettuce-specific DNA binding motifs.

We found 265 unique DAP-seq motifs which are significantly enriched (adjusted p<0.05 and enrichment ratio >2) in the gene promoters of at least 1 module (Supp Dataset 5d), using shuffled promoter sequences as the comparator. Every module had DAP-seq DNA binding motifs significantly enriched in its gene promoters. Supp Figure 7 shows a subset of the enriched motifs: the 6 most significantly enriched motifs for each module, the top 4 significantly enriched motifs which are unique to a module and the top 3 enriched motifs corresponding to an individual TF family. This includes 109 of the 265 significantly enriched DAP-seq motifs. Supp Figure 7 clearly shows that the ERF TFs are a dominant TF family, with many ERF DNA-binding motifs significantly enriched across both upregulated and downregulated modules. 17/20 modules were enriched for at least 1 ERF binding motif. In contrast, WRKY DNA-binding motifs showed enrichment in either a single or two modules (mostly in modules 1 and 2). As seen in Figure 4, module 1 is enriched for DEGs involved in phytohormone responses (JA, SA and ET) and here we see that it is also enriched in DNA-binding motifs of TFs known to mediate these responses. The DNA-binding motif of ERF1, a key regulator of the ET/JA response, is enriched in module 1 (271 input/94 shuffled). The DNA binding motif of WRKY33, another key JA regulator, is specifically enriched in module 1 (215 input/95 shuffled) as is the WRKY50 motif (140 input/ 53 shuffled). WRKY50 is known to bind the PR1 promoter, activating SA-responsive gene expression in an NPR1-independent manner (Hussain et al., 2018; Johnson et al., 2003). Lettuce orthologues of ERF1 and WRKY33 were also identified within module 1. This data indicates that we are able to identify DNA-binding motif elements which are highly conserved from Arabidopsis to lettuce and are likely facilitating differential gene expression in response to necrotrophic fungal infection.

**Figure 7:**
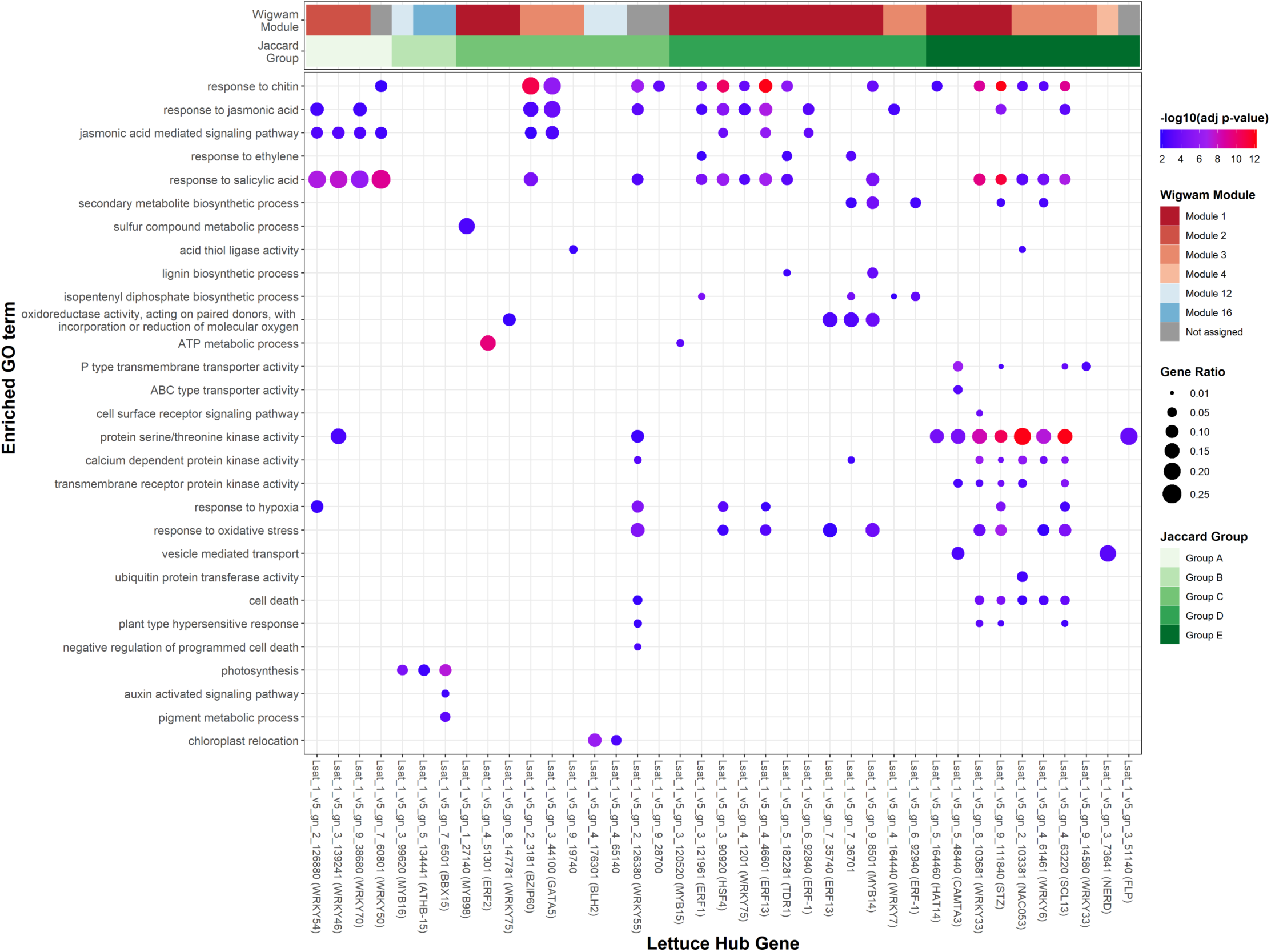
GO-term enrichment in GRN-predicted targets of Lettuce TFs. We perform Arabidopsis orthologs of the GRN-predicted target genes for each lettuce transcription factor. We have selected 29 key GO-terms for the heatmap, and shown all 39 lettuce hubs whose targets are significantly enriched (p-adjust <0.01) for at least 1 of the selected GO-terms. Colour of the point represents the statistical significance of the enriched (-log10 transformed adjusted p-value), with red dots showing higher significance. Size of the point represents the “GeneRatio”, number of predicted targets whose closest Arabidopsis orthologue is attributed with the GO-term as a proportion of the total number of predicted targets.

### A causal gene regulatory network predicts key transcriptional regulators of the lettuce response to *B. cinerea* and *S. sclerotiorum* infection

Co-expressed modules and promoter analysis above can predict regulatory interactions for experimental testing, however, the accuracy and confidence of regulatory predictions can be strengthened by the inclusion of additional data sets, and using network inference rather than single module approaches. To this end, we constructed a gene regulatory network (GRN) using four independent datasets: the time series data from lettuce inoculated with*B. cinerea*, the time series data from lettuce inoculated with *S. sclerotiorum* (Ransom et al., 2023) and single timepoint expression data from 21 diverse lettuce accessions following*B. cinerea* and *S. sclerotiorum* inoculation (Pink et al., 2022). We used the random forest OutPredict algorithm (Cirrone et al., 2020) to construct the GRN with the 4362 DEGs common to the *B. cinerea* and *S. sclerotiorum* infection time series as the input genes (Supp Dataset 5a). This includes 251 genes which were designated as TFs and hence are potential regulators of all other input genes. OutPredict tests potential regulator (in this case TF) expression profiles as predictors of the expression of all other input genes and outputs the likelihood of each TF influencing expression of each gene across all datasets.

The final GRN was constructed using the top 1% highest confidence TF-gene interactions and consisted of 3,382 genes (including 226 TFs) and 10,947 regulatory edges (Supp Dataset 6). The majority of TFs in the final network have a small influence on the expression of other genes in the network with 99 TFs (44%) having <10 predicted targets and 159 TFs (70%) <40 predicted targets. However, about a third of the TFs (hub genes) are predicted to have a very large influence on transcriptional reprogramming in response to necrotrophic fungal infection; 33 TFs (15%) have between 40 and 100 predicted targets and 34 TFs (15%) have ≥100 predicted targets (Figure 5A, Supp Dataset 7a).

The lettuce TFs predicted to have large numbers of downstream target genes include genes orthologous to Arabidopsis TFs known to impact defence against *B. cinerea*, such as WRKY33 (348 predicted targets)(Zheng et al., 2006), ERF1 (256 predicted targets)(Berrocal-Lobo et al., 2002), CAMTA3/SR1 (162 predicted targets)(Galon et al., 2008), MYC2 (98 predicted targets)(Lorenzo et al., 2004) and MYB108/BOS1 (48 predicted targets)(Mengiste et al., 2003; Cui et al., 2022)(Figure 5B). The prediction by the GRN of these known defence regulators having a significant impact on transcriptional reprogramming during pathogen infection, increases our confidence in the GRN to predict other (as yet unknown) regulators of the lettuce defence response against these two pathogens.

Analysis of the network node indegree distribution reveals that over a quarter of the genes in the network are predicted to be regulated by a single TF, however 2492 (74%) of the network genes are predicted to have multiple regulators (Figure 5C, D). Given this, we examined the extent to which TFs have shared downstream target genes by calculating the pairwise Jaccard Index (a measure of overlap) between predicted targets of the 67 hub TFs (≥40 outdegrees). This highlights five groups (A-E) of hub TFs which share predicted target genes (Figure 6). Each group contains TFs that are differentially expressed in the same direction during infection, with the exception of a single TF in group A and a single TF in group C. Group B contains only three TFs, all WRKY TFs in module 2, with a large overlap in predicted target genes with each other and almost no overlap in target genes with other hub TFs. *LsWRKY54* and *LsWRKY70* have a pairwise Jaccard index of 0.55, the highest overlap of any hub pair. Both these TFs are orthologues of key SA regulators, WRKY70 and WRKY54 (Zhang et al., 2010; Li et al., 2004), suggesting the presence of a distinct network for SA signalling.

Apart from this group of three WRKY TFs, the other groups of TFs with shared predicted target genes in the network contain TFs from different families. Group C is the largest but there is very little overlap between the target genes of each hub. Group D is dominated by TFs from module 1, with 6 ERF family TFs including orthologues of ERF1, ERF-1 and ERF13. This group also contains three WRKY TFs and two MYBs. Group E contains three additional WRKY TFs along with a NAC TF. This analysis therefore highlights the ability of the GRN to make distinct predictions for different members of the same TF family.

To assess the biological relevance of the specific predictions of target genes for a TF hub, we tested for GO term enrichment (using the Arabidopsis orthologues) in the genes uniquely predicted to be regulated by a TF hub, against a background of the genes expressed in the lettuce *B. cinerea* time series (Supp Dataset 7b). 255 GO terms were significantly enriched in the unique targets of at least 1 hub, and 48 (out of all 226) lettuce TFs had at least 1 enriched GO term in their unique GRN predicted targets. “Defence response to fungus” was significantly enriched in the targets of 26 TFs. Enrichment of selected GO terms and the corresponding 39 lettuce hubs is shown in Figure 7.

The value of the GRN above time series module analysis can be seen with the responses to JA and SA. Responses to these hormones (as judged by GO term enrichment) were both enriched in modules 1 and 2 (Fig 3), but the GRN predicts a group of 3 hubs (LsWRKY7, LsERF-1 and LsGATA5) that are predicted to specifically regulate JA responses; their unique targets are not enriched for response to SA (Figure 7).

We also see lettuce GRN hub targets being functionally enriched for GO terms that match the known function of Arabidopsis orthologues. For example, ERF1 is well-established as a key regulator of ethylene and jasmonate signalling in response to necrotrophic fungi (Berrocal-Lobo et al., 2002). The lettuce orthologue of ERF1, Lsat_1_v5_gn_3_121961 is a large network hub, and its target genes are significantly enriched for “response to jasmonic acid” and “response to ethylene” GO terms. WRKY70 is a key activator of the SA defence response (Zhang et al., 2010; Li et al., 2004). Its lettuce orthologue, Lsat_1_v5_gn_9_38680, has unique target genes (with an Arabidopsis orthologue) enriched for “response to salicylic acid” and “response to oomycetes”.

Lsat_1_v5_gn_2_103381 (LsNAC53) is a large GRN hub with unique target genes enriched for ”cell death” and “ubiquitin-protein transferase activity”, the only hub enriched for this term. LsNAC53 is a putative orthologue of AtNAC53 (also known as AtNTL4, NAC with transmembrane motif 1-like 4). AtNAC53 has been shown to regulate proteasome stress redundantly with its close homolog, NAC078 (Gladman et al., 2016). In addition, AtNAC53 is known to promote cell death and reactive oxygen species (ROS) production during drought stress, through directly activating expression of reactive burst oxidase homolog (RBOH)-encoding genes (Lee et al., 2012). In the GRN, a lettuce orthologue of *RBOHD* (*Lsat_1_v5_gn_5_9460*) is a predicted target gene of *LsNAC53*. *LsNAC53* and *LsRBOHD* show very similar expression patterns in both the *B. cinerea* and *S. sclerotiorum* inoculation time series (Figure 8) as well as correlation of expression with *LsNAC53* in the lettuce diversity panel transcriptome data, particularly after *S. sclerotiorum* infection (R=0.94). Orthologues of two other genes associated with cell death in Arabidopsis (*Necrotic spotted lesions 1 and 2, NSL1* and *NSL2*)(Noutoshi et al., 2006; Morita-Yamamuro et al., 2005)are also predicted targets of *LsNAC53* in the GRN and show a similar tight co-expression pattern with*LsNAC53* (Figure 8).

**Figure 8:**
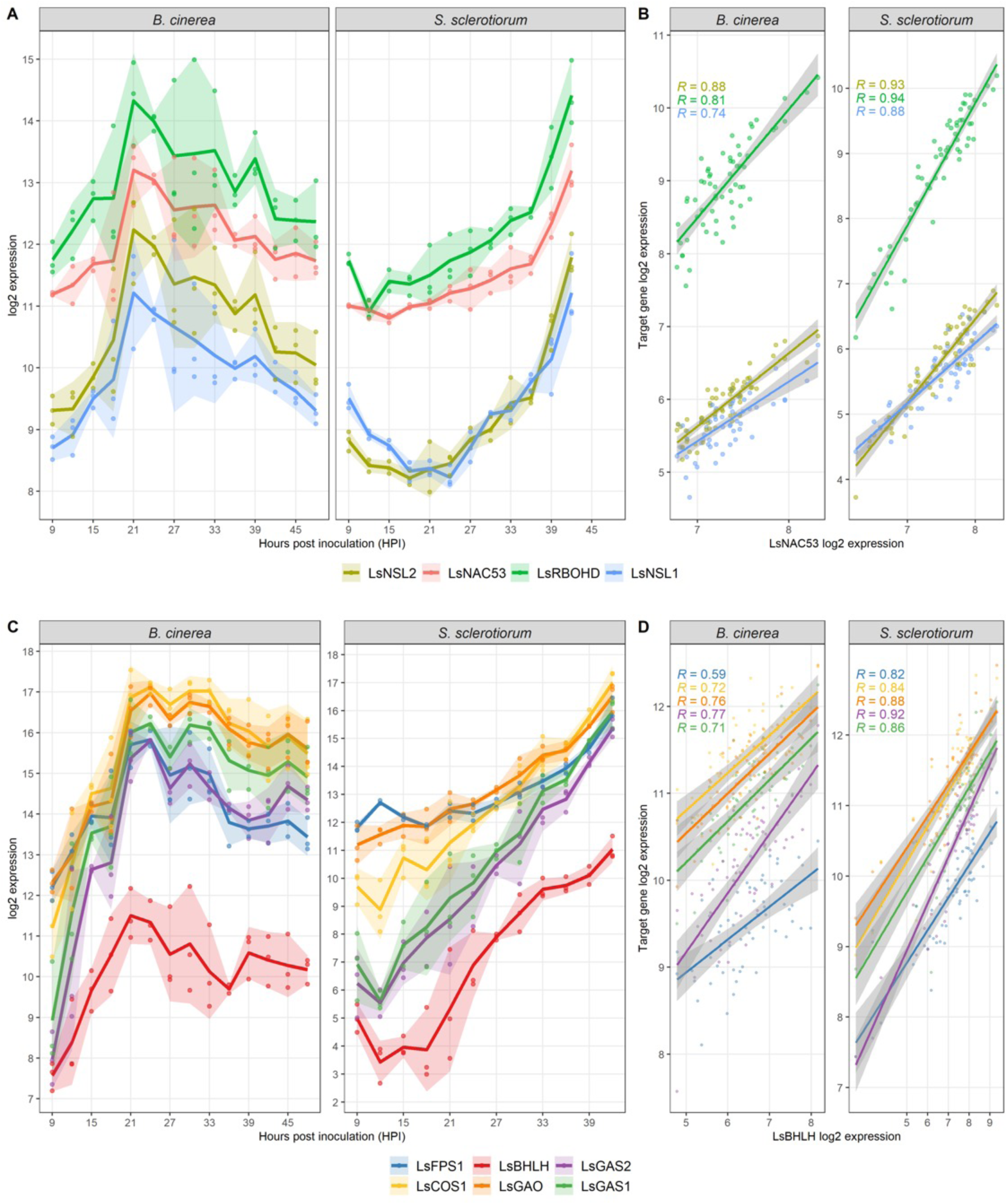
Expression profiles of the transcription factors *LsNAC53* and *LsBHLH* and their predicted downstream targets in the gene regulatory network. The expression of *LsNAC53* and its predicted targets *LsRBOHD*, *LsNSL1* and *LsNSL2* in lettuce following inoculation with *B. cinerea* (this study) and *S. sclerotiorum* (Ransom et al. 2023)(A) and the expression of the target genes compared to *LsNAC53* expression across 21 different lettuce accessions after pathogen inoculation (Pink et al. 2022) (B). C) and (A) D) show the expression profiles of LsBHLH and its predicted target genes (*LsFPS1*, *LsGAS1*, *LsGAS2*, *LsGAO*, *LsCOS1*) in the same data sets. *R* indicates the Pearson coefficient of correlation between expression of each target gene and its respective predicted regulator.

As mentioned above, lettuce synthesises a diverse range of sesquiterpene lactones (STLs), at least one of which has anti-fungal activity against *B. cinerea* in vitro (Bennett et al., 1994; Sessa et al., 2000b). In the GRN, multiple STL biosynthetic enzyme-encoding genes are predicted to be regulated by a single lettuce β helix-loop-helix TF (LsBHLH). These include genes encoding: a Farnesyl diphosphate synthase (*LsFPS1*), Germacrene A synthase (*LsGAS*), Germacrene A oxidase (*LsGAO*) and costunolide synthase (*LsCOS*). The expression profiles of *LsBHLH* and these predicted downstream targets are shown in Figure 8. This network prediction may not only identify a key transcriptional regulator of these specialised biosynthetic genes but also identify additional enzymes involved in the synthesis of this diverse family of compounds.

These examples highlight the ability of the lettuce GRN to not only predict TF hubs that impact disease resistance and associate these hubs with functional defence processes, but also to predict specific TF-target gene regulation that appears biologically relevant.

### Opposing functions of BOS1 in lettuce and Arabidopsis

We selected two hub TF genes from the network for functional testing: *Lsat_1_v5_gn_6_70301* (*LsBOS1*) with is orthologous to Arabidopsis *BOS1*, and *LsNAC53* highlighted above. The Arabidopsis *BOS1* gene encodes a MYB transcription factor (MYB108) that is upregulated during infection of Arabidopsis by *B. cinerea* (Windram et al., 2012; Mengiste et al., 2003). Despite the upregulation of this gene during infection, in Arabidopsis BOS1 has been shown to promote susceptibility to *B. cinerea* (Cui et al., 2022). The two putative lettuce orthologues of Arabidopsis *BOS1* (*Lsat_1_v5_gn_6_70301* and *Lsat_1_v5_gn_6_117600*) were significantly upregulated in response to both *B. cinerea* and *S. sclerotiorum* infection in lettuce (Supp Figure 8). Furthermore, both were hub genes in the GRN predicted to regulate 48 (Lsat_1_v5_gn_6_70301) and 39 (Lsat_1_v5_gn_6_70301) downstream target genes. To test whether the function of *BOS1* is conserved between Arabidopsis and lettuce, we generated transgenic Arabidopsis lines constitutively expressing *Lsat_1_v5_gn_6_70301* (named *LsBOS1* and selected due to the greater number of downstream target genes) under control of the 35S promoter. Two independent *p35S::LsBOS1* homozygous lines were selected from T2 lines showing Mendelian inheritance of the T-DNA selectable marker and were shown to express *LsBOS1* (Supp Figure 8). Both lines show increased resistance to *B. cinerea* with statistically significant reduced lesion size compared to wildtype Arabidopsis (Figure 9). This suggests that this lettuce*BOS1* gene (*Lsat_1_v5_gn_6_70301*) is acting as a positive regulator of plant defence against *B. cinerea*, in contrast to the Arabidopsis *BOS1* which promotes susceptibility to this pathogen. Despite opposing functions in defence, the altered disease resistance in these transgenic Arabidopsis lines demonstrates successful prediction of defence regulators by the OutPredict GRN.

**Figure 9:**
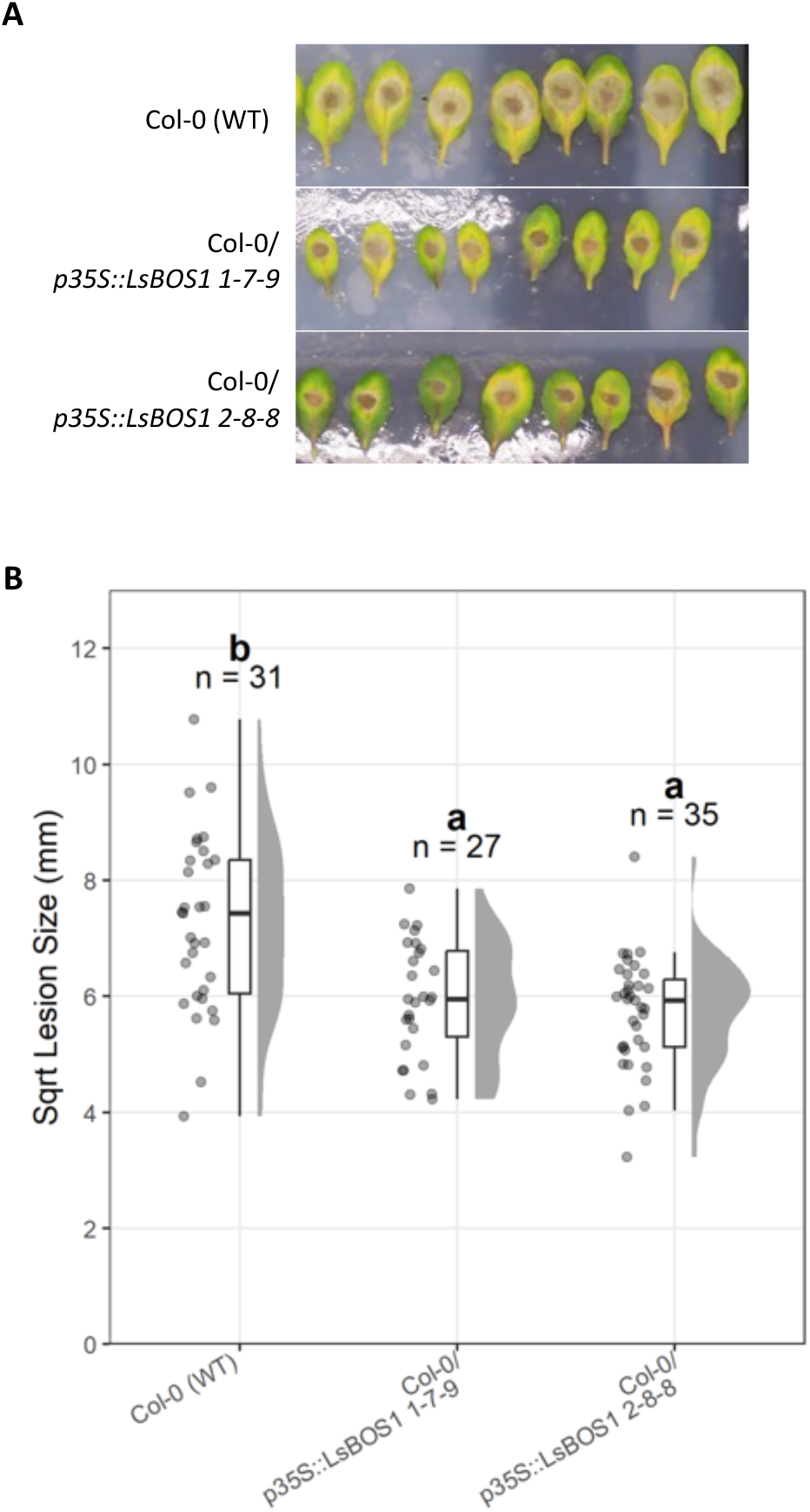
LsBOS1 acts as a positive regulator of *B. cinerea* resistance. (A) Representative images of Col-0 and two independent p35S::LsBOS1 transgenic lines (1-7-9 and 2-8-8) 72 hours post inoculation with *B. cinerea* “pepper” spores. Both transgenic lines exhibit stunted growth. **(B)** Quantification of (A) showing the square-root area of necrotrophic lesion, individual data points as well median (in box plot) and distributions. Letters represent statistical significance groupings – Tukey HSD p<0.05. N represents the number of lesion measured per genotype.

### GRN identification of a conserved novel defence regulator, NAC53

As outlined above, LsNAC53 is a putative orthologue of AtNAC53 (also known as AtNTL4). AtNAC53 regulates proteasome stress (redundantly with NAC078 [78]) and ROS production/cell death during drought stress, via RBOH gene expression (Lee et al., 2012). In addition to the conserved DNA binding domain both AtNAC53 and LsNAC53 have a C-terminal transmembrane domain, which in Arabidopsis has been shown to tether the TF to the plasma membrane (Kim et al., 2010). This prevents nuclear localisation and activity of AtNAC53 until the DNA binding domain is cleaved in response to drought, enabling it to move to the nucleus and activate gene expression.

Given the ROS/cell death promoting function of AtNAC53 we hypothesised that this TF would negatively impact plant resistance to necrotrophic fungal pathogens, despite expression of both *AtNAC53* and *LsNAC53* being upregulated following infection with *B. cinerea* (Windram et al., 2012)and this study) and, for LsNAC53, *S. sclerotiorum* (Ransom et al., 2023). We obtained seed of the previously characterised AtNAC53 T-DNA mutant, *nac53-1/ntl4-1* (Lee et al., 2012), and tested the susceptibility of this mutant line to *B. cinerea* using our detached leaf assay. Compared to wildtype Col-0 Arabidopsis, the*nac53-1* mutant showed increased resistance (smaller lesion size)(Figure 10A), suggesting AtNAC53 does indeed function as a negative regulator of *B. cinerea* defence.

**Figure 10:**
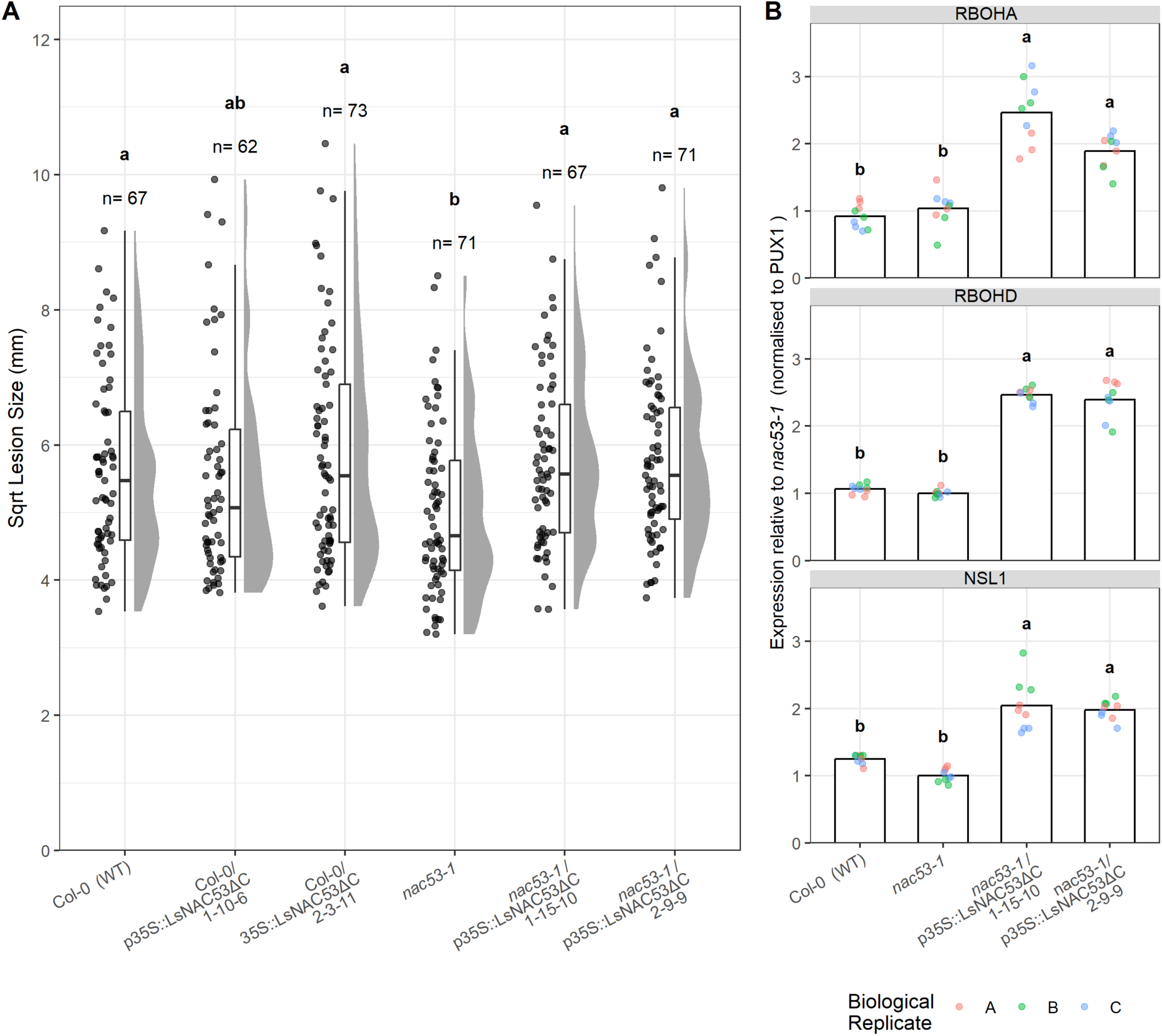
LsNAC53ΔC functionally complements *nac53-1* as a negative regulator of *B. cinerea* defence. (A) Lesion size after *B. cinerea* inoculated of detached leaves, 72 hours post inoculation. Arabidopsis genotypes are wildtype Col-0, *nac53-1* mutants, and constitutively expressed LsNAC53 (lacking the transmembrane domain) *p35S::LsNAC53ΔC* in both Col-0 and *nac53-1* backgrounds. Individual lesion sizes as well as the median and distribution of data points are shown. N = number of lesions measured, and letters indicate statistically significant differences as assessed by Tukey’s Honest difference. B) Expression of Arabidopsis genes RBOHA, RBOHD and NSL1 in wildtype Col-0, *nac53-1* mutant and transgenic Arabidopsis *nac53-1* mutants expressing truncated *LsNAC53* under control of the 35s promoter (*nac53-1 p35S::LsNAC53ΔC).* Three technical replicates of three biological replicates are shown with expression normalised to that of AtPUX1 and shown relative to expression in *nac53-1*. Letters indicate statistically significant differences based on Tukey’s Honest difference test.

To determine whether LsNAC53 is also a negative regulator of defence against*B. cinerea*, we generated transgenic Arabidopsis lines constitutively expressing *LsNAC53* without the transmembrane domain (p35S::LsNAC53ΔC) in both wildtype Col-0 and *nac53-1* mutant backgrounds. Two independent homozygous lines were selected in each genetic background which show similar levels of *LsNAC53* expression (Supp Fig 9).

Col-0 *p35S::LsNAC53ΔC* lines show no gain-of-function phenotype, with *B. cinerea* lesion size very similar to that of the wildtype Col-0. However, both independent*nac53-1 p35S::LsNAC53ΔC* lines have significantly larger *B. cinerea* lesions than *nac53-1* (Fig 10A), demonstrating that expression of the lettuce NAC53 TF can functionally complement the lack of AtNAC53. This suggests that LsNAC53 is a functional orthologue of AtNAC53, and also acts as a negative regulator of *B. cinerea* defence.

As outlined above, Arabidopsis NAC53 TF activates expression of the*RBOH* genes *A*, *C* and *E* (Lee et al., 2012) and in the GRN, *LsNAC53* is predicted to regulate *LsRBOHD*, *NSL1* and *NSL2* (Figure 8). Analysis of gene expression in these transgenic Arabidopsis lines demonstrated that LsNAC53 can activate expression of two Arabidopsis *RBOH* genes (*A* and *D*) as well as Arabidopsis *NSL1* expression (Figure 10). As seen before, the *nac53-1* mutant did not reduce expression of these genes under non-stressed conditions, but clear induction of expression was seen in the presence of the truncated *LsNAC53*, validating these GRN predictions.

## Discussion

Here we present a high-density time series transcriptome dataset capturing gene expression after *B. cinerea* and mock inoculation of lettuce leaves. Comparing the response to a similar time series dataset following *S. sclerotiorum* infection, revealed a core set of 4362 lettuce genes that change in expression (in the same direction) in response to infection by both pathogens. An earlier lettuce -*B. cinerea* RNAseq dataset was published with just three time-points (12, 24 and 48 hours post inoculation), identifying 1, 139 and 4598 DEGs at each respective time point (Cremer et al., 2013). In contrast, the time series presented in this paper has 14 time points, one every 3 hours, and has captured significant gene expression changes from 9 hpi. As seen during *B. cinerea* infection of Arabidopsis (Windram et al., 2012) the majority of gene expression changes occur before significant growth of the pathogen (Fig 2) or lesion development (Fig 1). A significant proportion of the lettuce genes differentially expressed in response to *B. cinerea* are orthologous to DEGs in Arabidopsis (Supp Fig 4) although there are some clear differences. Plant defensins play a major role in Arabidopsis defence against *B. cinerea* with several dramatically upregulated during infection (Windram et al., 2012; Ingle et al., 2015), but in lettuce these genes are generally not changing in expression with one member of the family downregulated (Supp Fig 6). In addition, the Arabidopsis BOS1 TF is a negative regulator of defence against*B. cinerea* (Mengiste et al., 2003; Cui et al., 2022) whereas the lettuce gene (when expressed in Arabidopsis) appears to be a positive regulator (Fig 9). Interestingly the cotton orthologue of BOS1 (GhMYB108) is a positive regulator of resistance to *Verticillium dahlia* and *B. cinerea* (Cheng et al., 2016). These examples show the importance and value of analysing defence responses (even to the same pathogen) in different species.

The high resolution of our time series data provides insight into the timing and sequence of pathogen-induced transcriptional reprogramming. We used Wigwams (Polanski et al., 2014) to identify modules of genes that are co-expressed in response to *B. cinerea* and *S. sclerotiorum* to reduce the complexity of the data and identify groups of genes with a shared function that are similarly regulated. This did highlight known defence responses (such as JA and ET signaling, receptor signaling) as well as lettuce-specific processes such as the synthesis of sesquiterpene lactones. However, some of these modules are very large. We still observe the majority of DEGs changing in expression during a short window (21-24 hpi) and finer time points in this region could help in separating gene expression profiles (and biological processes) further. However, unlike application of hormones or defence elicitors (Hickman et al., 2017; Bjornson et al., 2021) where responses occur within minutes, the requirement for spore germination and growth of the pathogen, and the impact of the environment on this, means the first transcriptional responses can only be detected hours after inoculation and timing can vary between experiments making it hard to accurately predict the critical window for analysing more time points.

However, the availability of high-resolution time series data enables the inference of a causal GRN model of the regulatory events underlying transcriptional reprogramming during infection. The power of such network inference is increased with the availability of two such time series (lettuce inoculated with *B. cinerea* and *S. sclerotiorum*). Furthermore, the ability of the OutPredict algorithm to combine time series and static (single time point) data meant that we could also incorporate transcriptome data from 21 different lettuce accessions after infection with *B. cinerea* and *S. sclerotiorum* (Pink et al., 2022). Time series data provides information on the relative timing of a TF and target gene, whereas the diversity set data provides an independent set of data highlighting correlation in expression between the TF and target genes. Combining these different types of data has likely increased the power of OutPredict and the accuracy of the resulting GRN.

The GRN we have generated (using the top 1% high confidence edges) not only predicts key regulators but is also able to predict downstream target genes of these regulators. Our confidence in the network model comes from i) the identification as hubs of orthologues of known Arabidopsis TFs that impact *B. cinerea* disease resistance (Fig 5); ii) the demonstration that LsBOS1 and LsNAC3 impact resistance to *B. cinerea* when expressed in Arabidopsis (Fig 9, 10); and iii) LsNAC53 is able to upregulate orthologues of its predicted target genes when expressed in Arabidopsis (Fig 8, 10). This GRN will advance our understanding of the transcriptional defence response in lettuce by identifying key regulators for experimental testing (with resulting data able to be used to improve the model), and highlighting the network topology, network motifs and cross-talk between different signaling pathways that is driving the ultimate defence response. Future work will aim at validating the GRN in lettuce and developing GRN models with the ability to simulate the impacts of network perturbation not just on expression of GRN genes, but on disease resistance against these important pathogens.

## Methods

### Pathogen inoculation time series experiment

Lettuce cv. Saladin were grown for 4 weeks in Levington’s M2 soil in the greenhouse at approx. 18°C with day length supplemented to 16 hours. The third leaf from each plant was removed and placed on 0.8% agar in 35 x 23 cm propagator trays. Leaves were inoculated with four 10 µL droplets of 5 x 10^5^ mL *B. cinerea* ‘pepper’ isolate (Windram et al., 2012) spore suspension in 50% (w/v) potato dextrose broth (PDB), 1% (w/v) guar, or mock inoculated with four 10 µL droplets of 50% PDB, 1% guar. B. cinerea spore suspensions were prepared as in (Pink et al., 2022). Inoculations were carried out halfway through the 16-hour light period. Lidded trays were placed in a controlled environment chamber under 16-hour light: 8-hour dark, 22^0^C at 95% humidity. A 1 cm corkborer was used to harvest a leaf disc surrounding each inoculation droplet, with the four discs from one leaf pooled and flash-frozen in liquid nitrogen. Three leaves (each from separate plants) were harvested for mock- and *B cinerea*-inoculations at each of the 14 time-points: 3-hour intervals from 9 to 48 hours post inoculation (84 samples in total).

### Gene expression profiling

RNA was extracted using Trizol (Thermo Fisher Scientific) with a lithium chloride purification. Sequencing libraries were prepared using the Illumina TruSeq RNA V2 kit and sequenced on a HiSeq 3000 generating 75 bp paired-end reads at the Wellcome Trust Human Genetics Centre. Read quality was checked with FastQC (Andrews, 2010). Raw reads were trimmed with trimmomatic (Bolger et al., 2014) and aligned to a combined*L. sativa* cv. Saladin – *B. cinerea* transcriptome using STAR (Dobin et al., 2013), achieving a median alignment of 90% (Supp Dataset 1). Transcript abundances were calculated with RSEM (Li and Dewey, 2011). We applied a low expression filter, keeping genes >1 count per million in at least 3 samples. Principal component analysis was performed using ‘prcomp‘ R function.

Pairwise differential expression analysis between mock and *B. cinerea* inoculated samples at each time-point was performed using the Limma-voom pipeline (Ritchie et al., 2015; Law et al., 2014). P-values for each time-point were combined using the Simes method (SIMES, 1986; Sarkar and Chang, 1997) obtaining a single combined p-value per gene for the time series, these were subsequently adjusted using Bonferroni-Hochberg (BH) (Benjamini and Hochberg, 1995) to account for multiple testing. Genes with a final adjusted p-value <0.01 were considered differentially expressed. The Time of First Differential Expression (TOFDE) was determined by examining the pairwise time-point comparisons. Genes with BH-adjusted p-values < 0.01 were considered differentially expressed at a specific time-point.

TOFDE was determined for each gene that exhibited differential expression across the entire time-series, by the earliest time-point at which they were identified as differentially expressed.

### Wigwams modules

The Wigwams algorithm (Polanski et al., 2014) (github.com/cyversewarwick/wigwams) was used to identify co-expressed modules within the set of DEGs common to both*B. cinerea* and *S. sclerotiorum* infection. The following parameters were used: ‘SizeThresholds‘ = 30, ‘Merging_Overlap‘ =0.82, ‘Merging_CorrelationFilter‘ = 0.89, ‘Mining_CorrelationNet =0.5, ‘Merging_MeanCorrelation‘ =0.93.

### Functional enrichment

Gene-ontology GO term enrichment analysis was performed using ‘enrichGÒ function within the clusterProfiler R package (Wu et al., 2021), performed with a significance threshold of p<0.05; p-values were corrected for multiple testing using Bonferroni-Hochberg. Arabidopsis GO term annotation was used and enrichment performed using the closest Arabidopsis orthologue (Reyes-Chin-Wo et al., 2017) of lettuce genes in the test set, against a background of all Arabidopsis genes with an identified lettuce orthologue expressed in the *B. cinerea* time-series.

Protein domain enrichment was performed using previously published InterProScan annotations (Reyes-Chin-Wo et al., 2017). A hypergeometric test was performed with the ‘phyper‘ R function using all genes expressed in the lettuce *B. cinerea* time series as the background. P-values were corrected for multiple testing using Bonferroni-Hochberg.

### Transcription factor binding motif enrichment

Lettuce promoter sequences 1000 bp upstream from the predicted transcription start site (TSS) were extracted from the *L. sativa* cv. Saladin V8 genome (Reyes-Chin-Wo et al., 2017). The enrichment of Arabidopsis DAP-seq transcription factor binding sites (O’Malley et al., 2016) was tested within these lettuce promoter sequences using SEA (Simple Enrichment Analysis) within MEME-suite (Bailey et al., 2009; Bailey and Grant, 2021). Shuffled input sequences (1000 bp lettuce promoters) were used as background.

### Gene Regulatory Network

A gene regulatory network was constructed with OutPredict (Cirrone et al., 2020) with genes differentially expressed in both *B. cinerea* and *S. sclerotiorum* time series in the same direction used as input. 251 of these DEGs, identified as TFs using the PlantTFDB predictor [93], were identified as potential regulators. The GRN was trained on expression of these genes in two pathogen infection time series (*B. cinerea* – this study, *S. sclerotiorum* – (Ransom et al., 2023) and two previously published lettuce diversity panel data sets (Pink et al., 2022). All expression data was scaled to ensure comparability between experiments. The OutPredict random-forest model was trained with 300 estimators and a test-train split ratio of 0.15. The model was trained on both the time series and single time point data, with the leave-out test set from the time-series. OutPredict calculates an Importance score for the influence of each of the 251 TFs on every non-self-target gene. The top 1% highest confidence edges were included in the final network, comprising 3,382 nodes (including 226 TFs) and 10,947 edges. Pairwise Jaccard-index (TF1-TF2 target intersection/TF1-TF2 target union) was calculated to quantify predicted target overlap of TFs.

### Transgenic Arabidopsis lines

The *nac53-1* (SALK_009578C) Arabidopsis mutant line was obtained from NASC. Lettuce *BOS1* (*Lsat_1_v5_gn_6_70301)* and *NAC53* (Lsat_1_v5_gn_2_103381) sequences were amplified from *L. sativa* cv. Saladin cDNA generated from mRNA extraction of *B. cinerea* infected leaf material. Primers with attB1 and attB2 extensions (Supplementary Dataset 8) were used to amplify the full-length coding sequence of LsBOS1 and a truncated version of LsNAC53 which lacked the C terminal transmembrane domain. Sequences were cloned and verified in the pDONR-Zeo vector, before cloning into the destination vector, pB2GW7 (Karimi et al., 2002) containing a 35S promoter. Stable Arabidopsis transformants were generated using the floral-dip method (Clough and Bent, 1998). Multiple independent homozygous transformed lines were selected, and expression of the transgene determined via qPCR of T_3_ homozygous plants. T_3_ lines were subsequently used for pathogen infection assays and analysis of downstream target genes.

### Arabidopsis-B. cinerea susceptibility assay

Arabidopsis-*B. cinerea* infection assays were performed as previously described (Windram et al., 2012). In summary, Arabidopsis plants were grown in P24 trays on Levington’s F2+Sand soil in controlled environment growth chambers (16 hour day length, 22°C day and night, 60% relative humidity) for 4 weeks. A single leaf was detached from a plant and placed in propagator trays containing a layer of 0.8% agar. Detached leaves were inoculated with 10 µL of *B. cinerea* spore suspension (pepper isolate) at a concentration of 1 x 10^5^ spores/mL, diluted in 50% filter-sterilised grape juice. Trays are sealed and placed back in the growth chamber at a relative humidity of 90%. Photographs are taken of the developing lesions at 72 hpi and lesion size measured using ImageJ software. Statistical differences between genotypes was determined using Tukey HSD test p<0.05 (Tukey, 1949).

### qPCR expression analysis

Arabidopsis seedlings were grown on ½ strength Murashige and Skoog (MS) media agar plates for 10 days under a 16 hr photoperiod. RNA extractions were performed from whole seedlings using Qiagen RNeasy Plant columns, with an on-column DNase digestion step. cDNA synthesis was performed using SuperScript III (Invitrogen), qPCRs with SYBR green. All qPCRs are performed with three technical replicates of three biological replicates (separate seedling pools). *PUX1* (*At3g27310*) or *PP2AA3* (*At1g13320*) was used as an internal control to normalise expression. Delta Ct (2−^ΔCt^) was used to analyse transgene expression level relative to an endogenous gene and delta-delta Ct (2−^ΔΔCt^) to calculate relative expression of an endogenous Arabidopsis gene.

### Data Analysis and Visualisation

Statistical analysis, data analysis and data visualisation were performed using R, unless stated otherwise. The “Tidyverse” collection of R packages was used for all data analysis and data manipulation (Wickham et al., 2019). Heatmap figures were visualised with ComplexHeatmap R package (Gu et al., 2016). Phylogenetic trees were generated using MEGA X (Kumar et al., 2018) and visualised with treeio (Wang et al., 2019) and ggtree R packages (Yu et al., 2016).

Details of specific lettuce genes named and discussed in this work are given in Supplementary Dataset 9.

## Author Contribution Statement

The study was conceived and designed by KD, JC, CY, FG, PH and DP with input from HP. Experimental work was carried out by AT, GH, OC, RL and HP with data analysis carried out by HP, RC, AT and RH and further data interpretation performed by HP and KD. The manuscript was written by HP and KD with input from all authors. All authors have approved the submission of this manuscript.

## Data Availability

All data is available either within the supporting information of this manuscript or in the NCBI Short Read archive under Bioproject PRJNA804213 (diversity set RNAseq data) and Bioproject PRJNA808232 (time series).

---

**Supp Figure 1:**
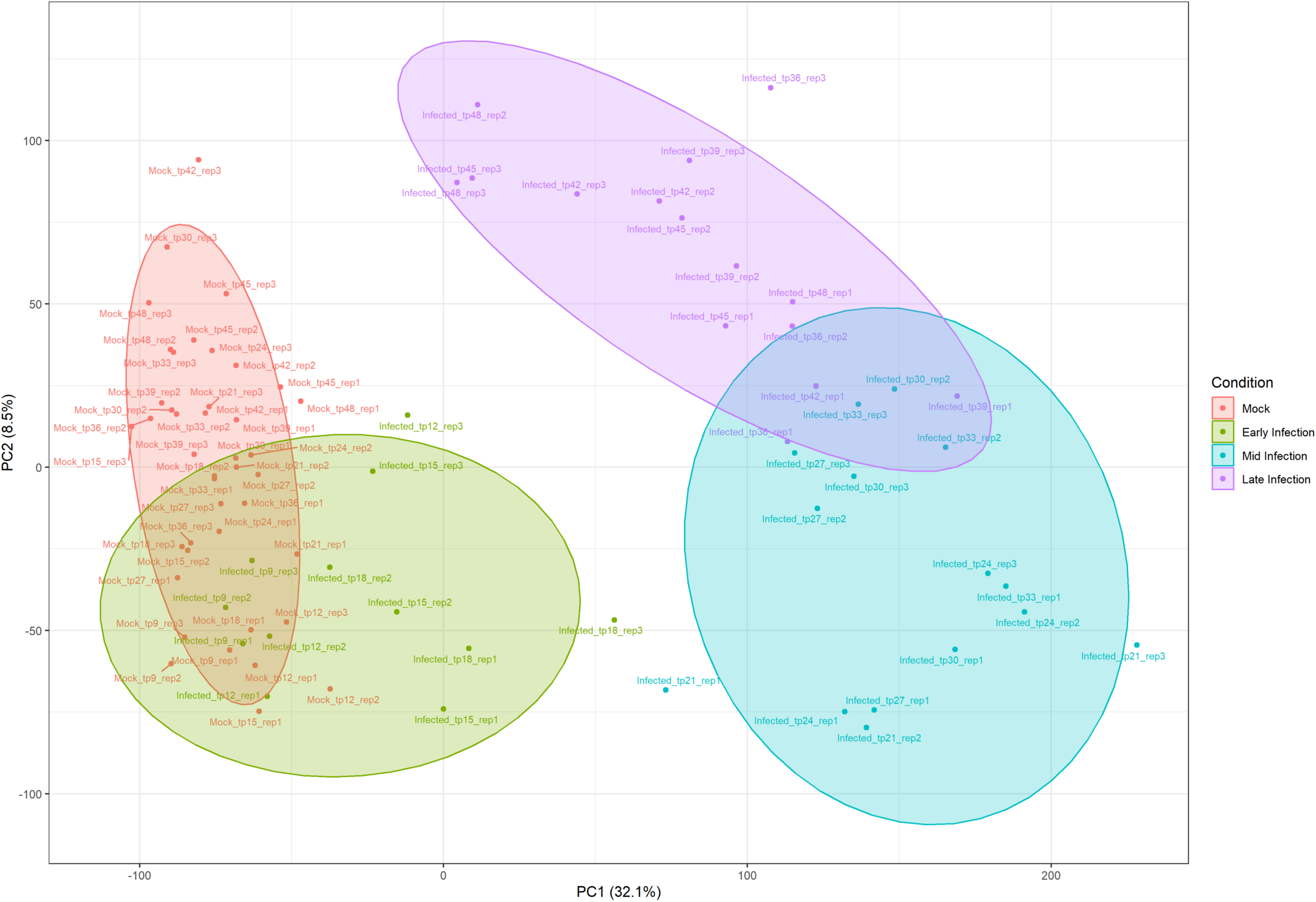
Principal component analysis of the lettuce gene expression data (TPM) demonstrates variability between samples is reflected by time point after inoculation. Mock-inoculated samples (red), early infection time points (9-18 hpi, green), mid infection time points (21-33 hpi, blue) and late infection time points (36-48 hpi, purple) are indicated. The proportion of variance explained by the first and second principal components is 32% and 8.5% respectively. Ellipses representing 90% confidence intervals around each group’s data points.

**Supp Figure 2:**
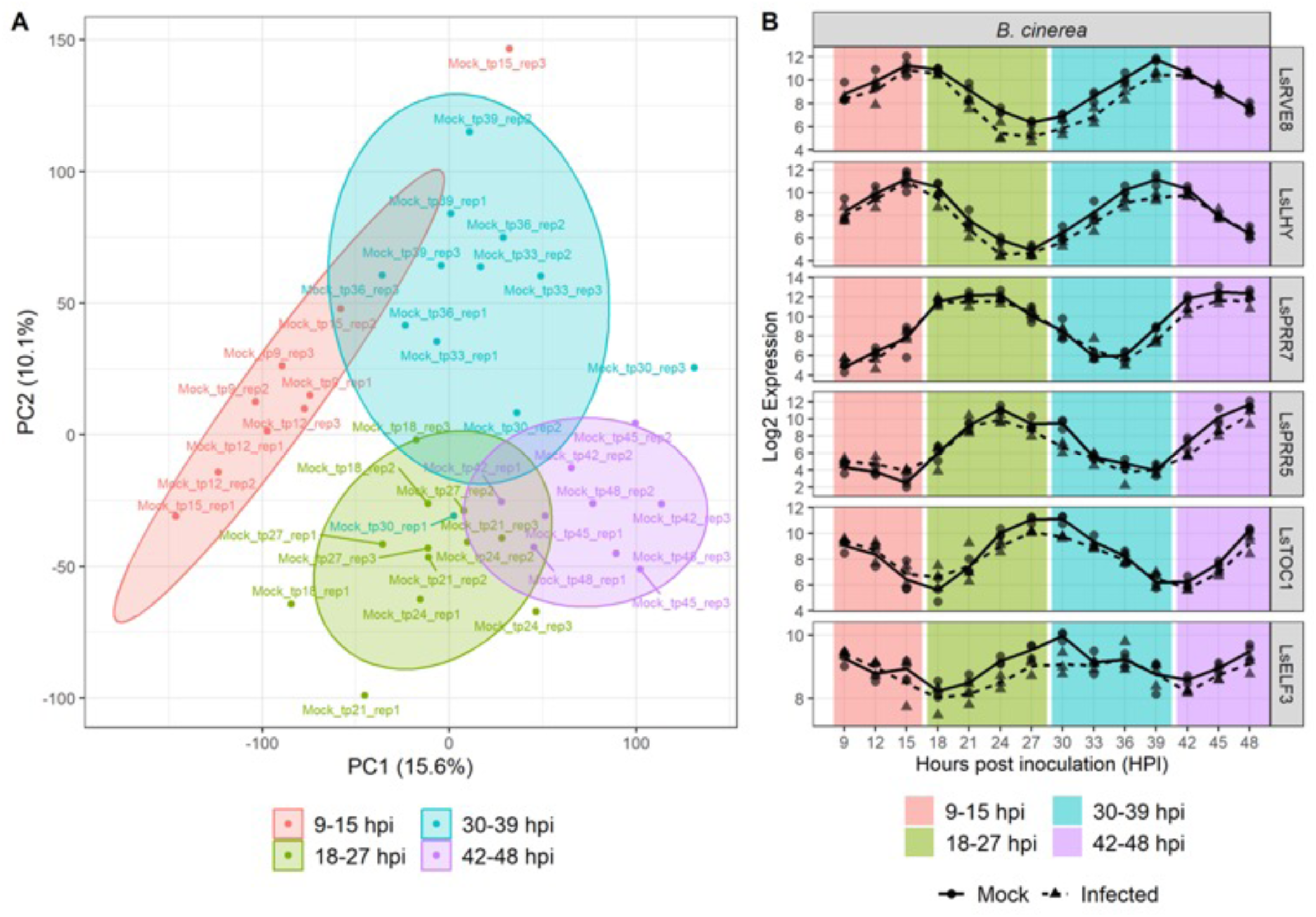
Circadian Oscillation observed in Lettuce time-series. **(A)** Principal Component Analysis of mock inoculated gene expression, PC1 is plotted on the X-axis, PC2 is plotted on the y-axis accounting for 15.6% and 10.1 &% of variation respectively. Samples between 9 and 15 hours post inoculation (hpi) are coloured in red, 18 to 27 hpi samples are green, 30 to 39 hpi in blue and 42 to 48 hpi. Lettuce orthologues of known circadian oscillators show rhythmic expression profiles in the time-series data., demonstrating that a single time-point 0 mock would not be a sufficient control across the time-series. Lsat_1_v5_gn_6_101520 = LsGI (Gigantea), Lsat_1_v5_gn_3_140720 = LsLHY (Late=elongated Hypocotyl), Lsat_1_v5_gn_8_4341 = LsPRR5 (Pseudo-response regulator 5), Lsat_1_v5_gn_2_115441 (LsPRR7) (Pseudo-response regulator 7), Lsat_1_v5_gn_2_109801 = LsRVE8 (Reveille 8) and Lsat_1_v5_gn_1_14320/Lsat_1_v5_gn_5_21481 = LsTOC1A/LsTOC1B respectively (Timing of CAB expression 1)

**Supp Figure 3:**
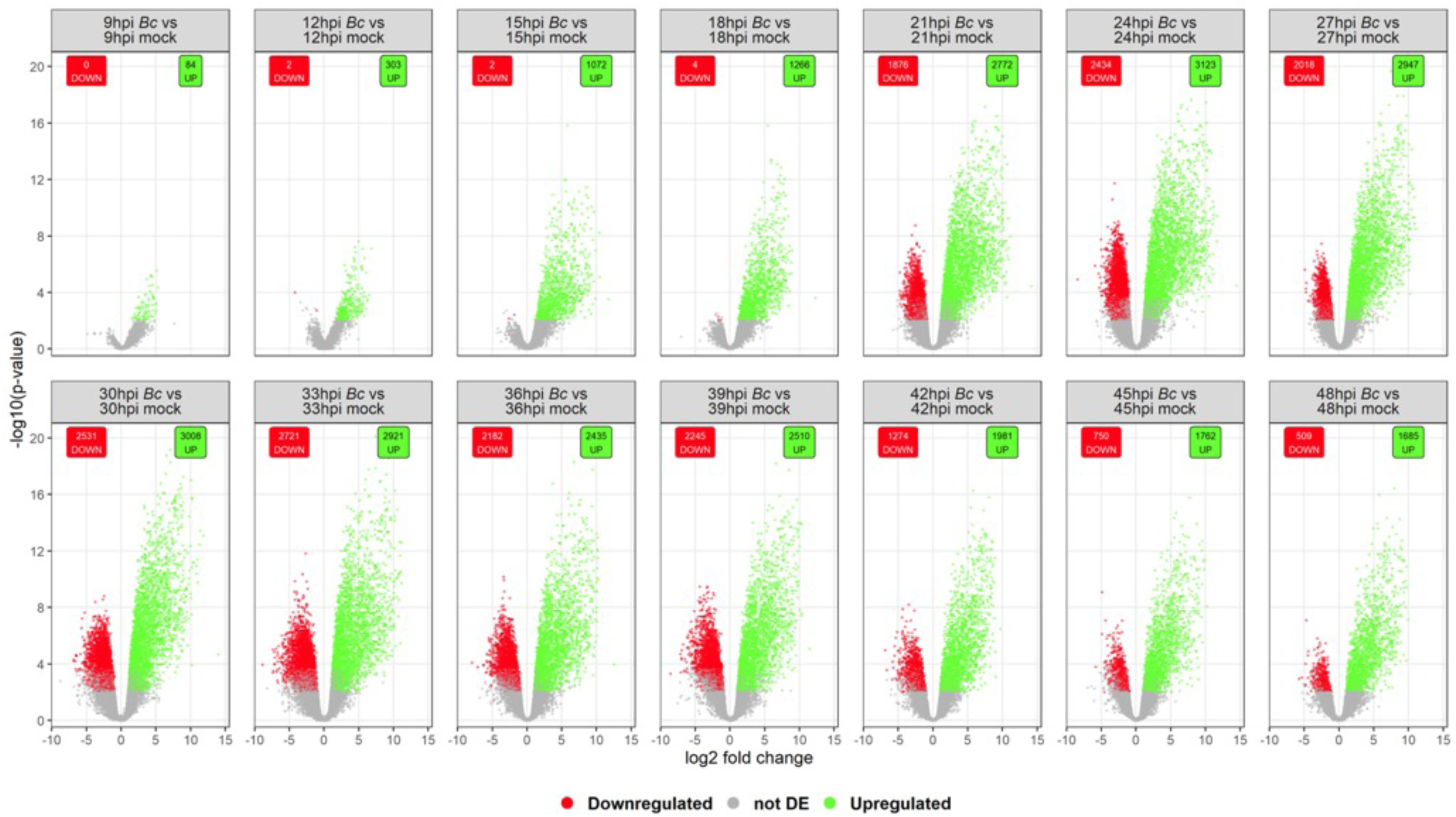
Volcano plots of *B. cinerea* vs mock inoculated at each individual time-point highlighting the scale of differential gene expression. Log_2_ fold-change is displayed on the X axis, and –log10 transformation of Bonferroni-Hochberg adjusted p-values (for the single time-point) are shown on the y axis. Significantly upregulated genes (B. cinerea inoculated versus mock inoculated samples) are highlighted in green, and significantly downregulated genes in red.

**Supp Figure 4:**
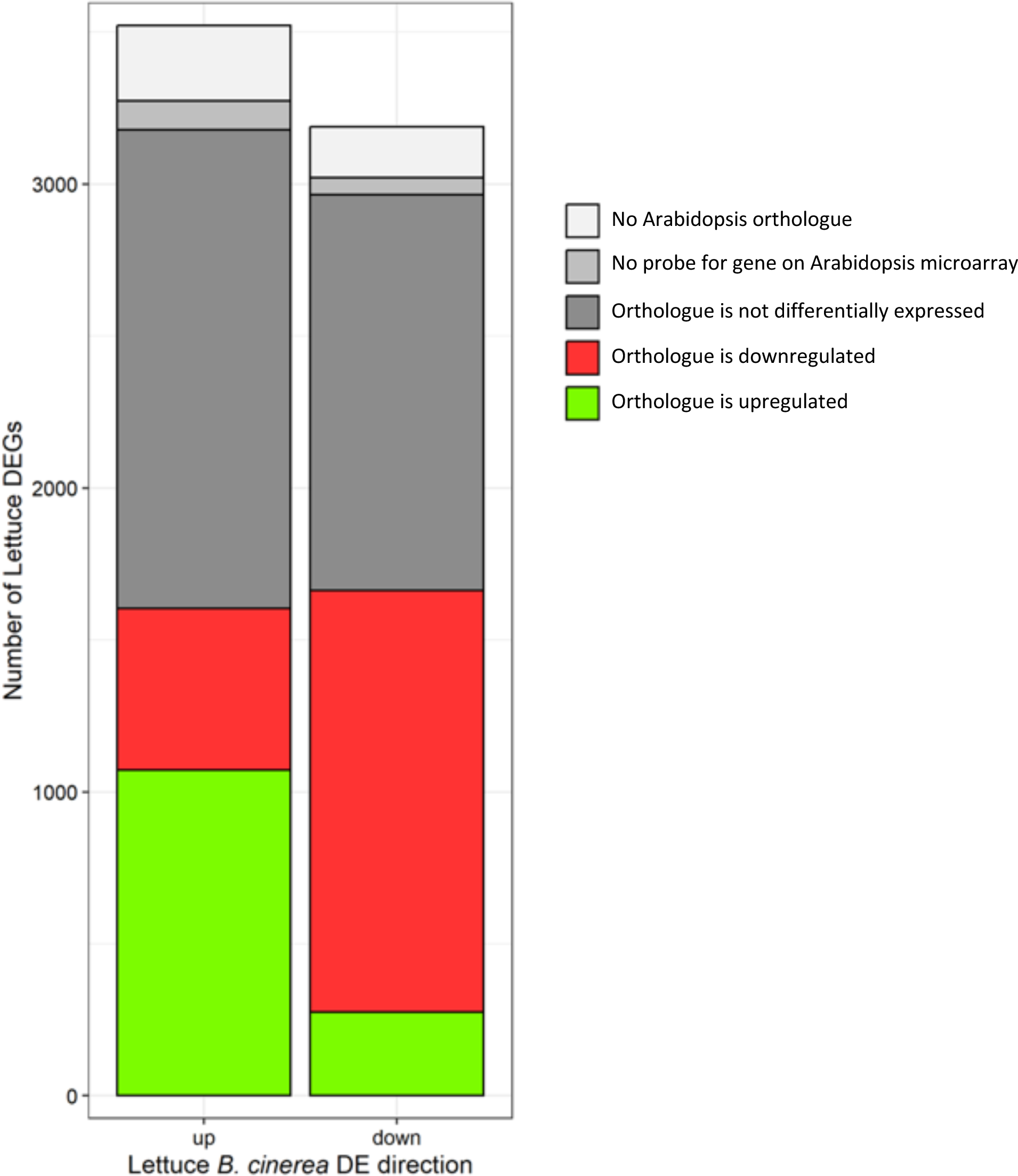
Species specificity of transcriptional reprogramming during *B. cinerea* infection. Bar chart showing the number of lettuce genes up and downregulated after inoculation with *B. cinerea*. Bars are coloured by the direction of differential expression following *B. cinerea* inoculation of the single closest Arabidopsis orthologue for each lettuce DEG (Windram el al, 2012). Orthologues were as in Reyes Chin-Wo et al. (2017).

**Supp Fig 5:**
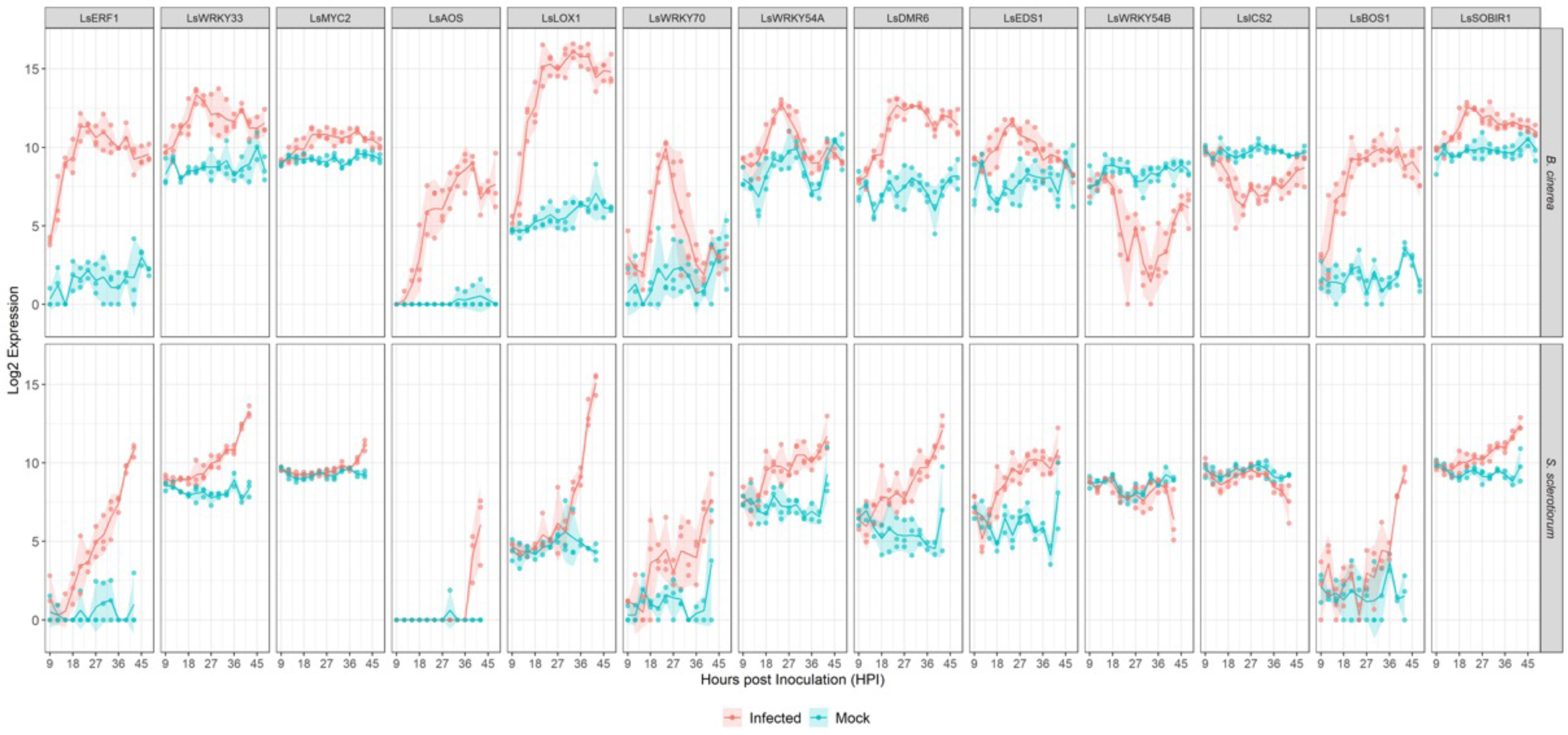
Lettuce orthologues of known Arabidopsis defence regulators are differentially expressed during both *B. cinerea* and *S. sclerotiorum* infection. Individual data points (log2 expression) are shown along with the mean and 95% confidence interval. Red indicates pathogen inoculated samples, blue are mock inoculated. Gene IDs listed in Supp Dataset X

**Supp Figure 6:**
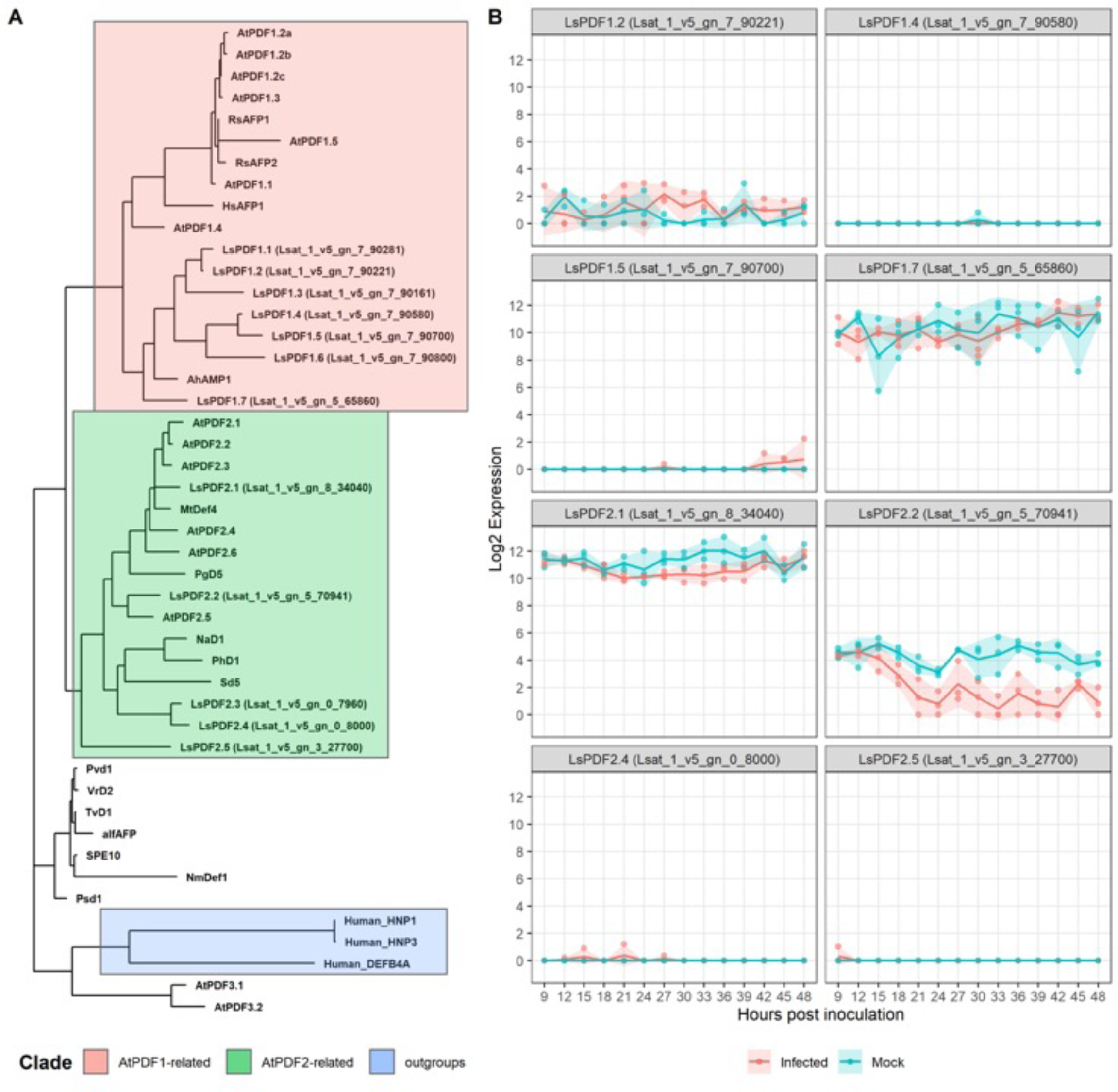
Phylogeny and expression profile during *B. cinerea* infection of lettuce defensins. A) A 750 bootstrap maximum likelihood phylogenetic tree of 13 putative lettuce defensins, Arabidopsis Plant Defensins (AtPDFs) and characterised anti-fungal defensins from other species (Lacerda et al, 2014). B) *B. cinerea* and mock inoculated time-series expression profiles of putative LsPDFs. Only 8 of the 13 LsPDFs were detected in at least 1 sample, with only 4 were consistently detected across the time series. Only 1 was differentially expressed, *Ls_1_v5_gn_5_70941*, which was downregulated.

**Supplementary Figure 7.**
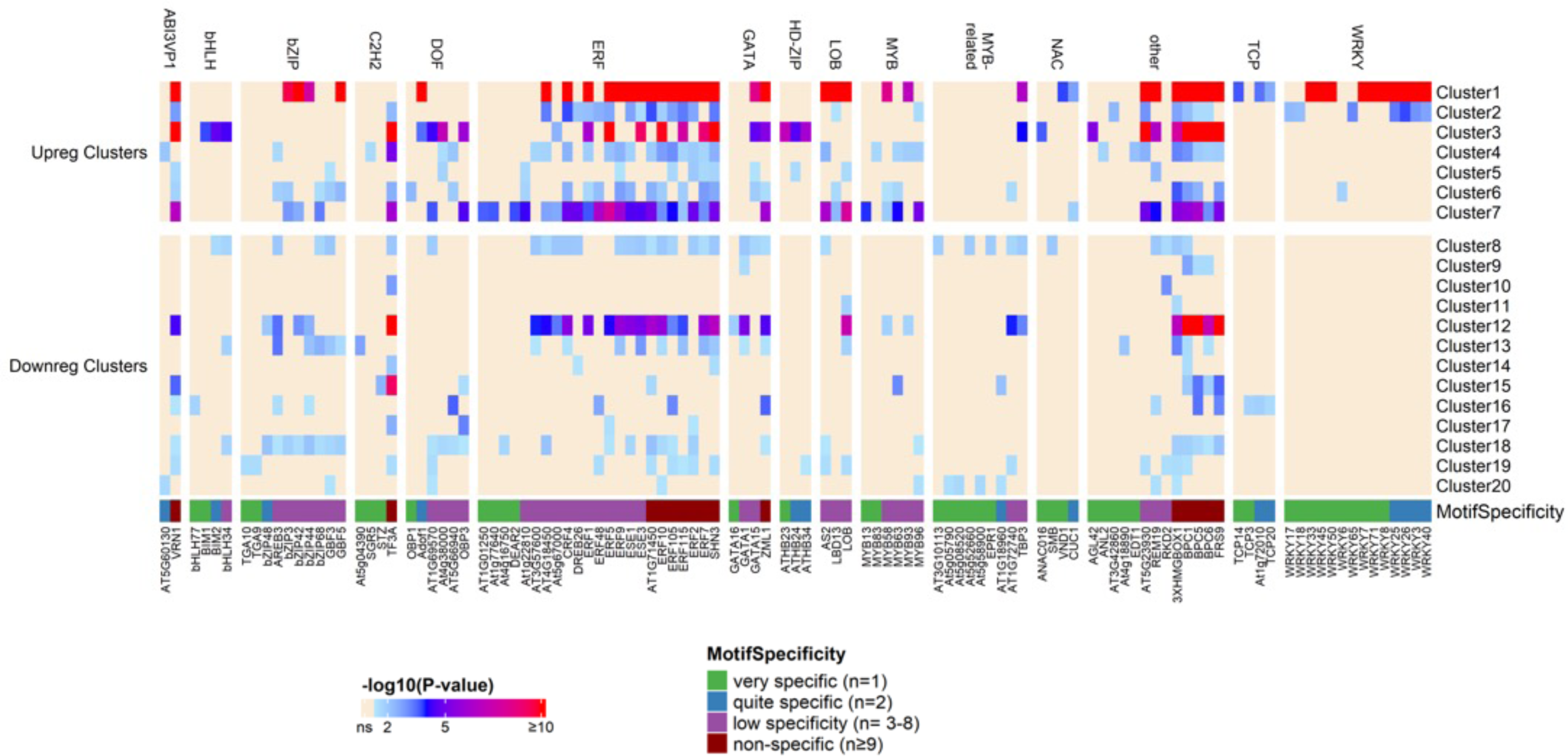
Enrichment of Arabidopsis DAP-seq DNA-binding motifs in 1 Kb promoters of lettuce DEGs. Columns indicate individual motifs, which are named according to the binding TF and grouped by their respective TF family. This heatmap shows 109/265 enriched motifs. We selected motifs which were either in the top 6 enriched motifs in a module, the top 4 enriched motifs which are unique to a single module or in the top 3 enriched motifs for an individual TF family. Rows represent the time-series modules. Colour indicates the significance of the enrichment (-log10 transformation of the adjusted p value) for a specific motif in a specific module, with white = non-significant (ns) enrichment. Significant enrichment is defined as p-adjust <0.05 and enrichment ratio >2. “Motif Specificity” is indicated in four classes, corresponding to the number of modules a motif is enriched in.

**Supplementary Figure 8:**
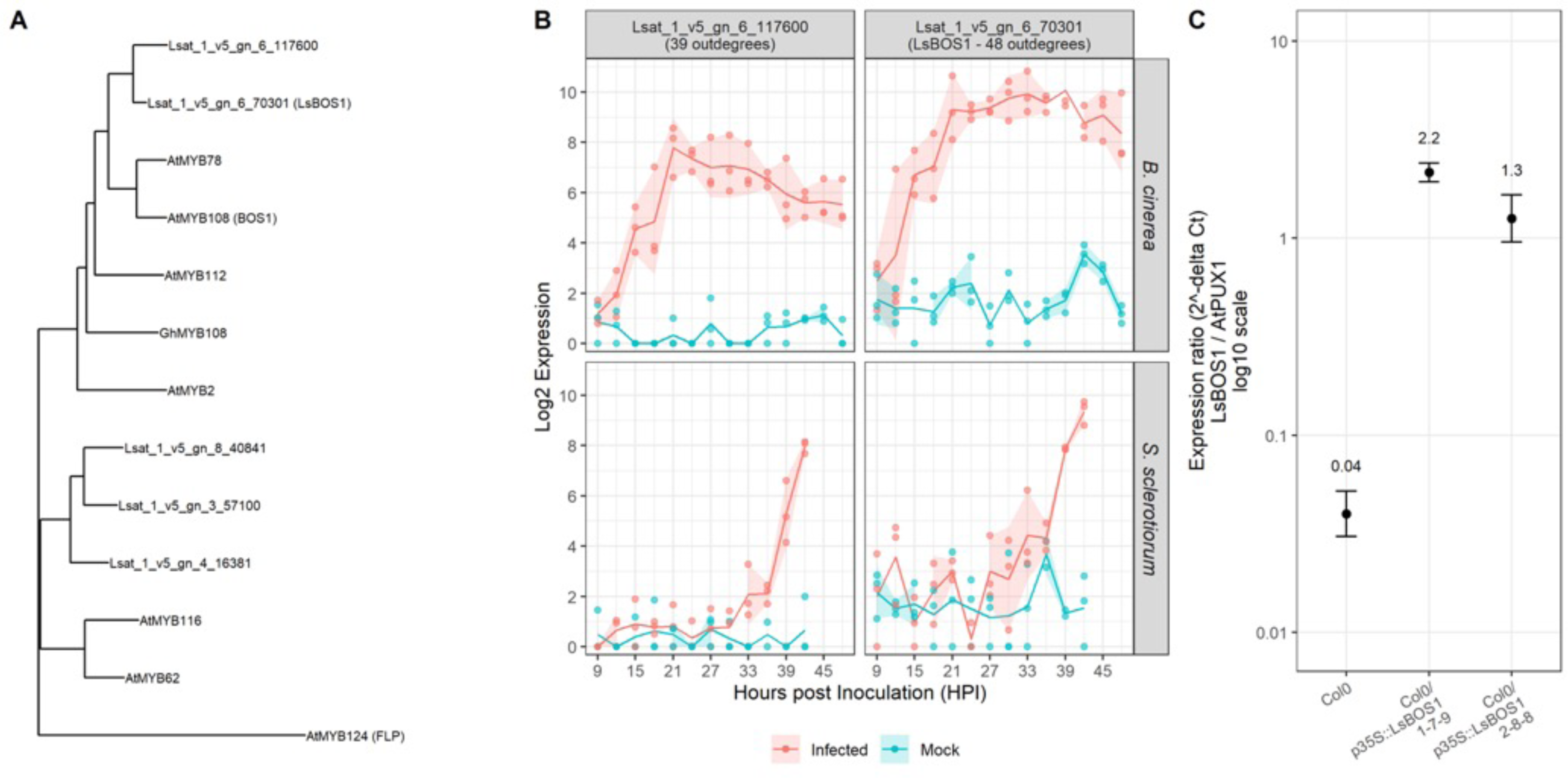
Data underlying experimental testing of LsBOS1. **(A)** A 3000-boostrap maximum likelihood phylogenetic tree of Arabidopsis MYB subgroup 20 (MYB2, MYB62, MYB78, MYB108 (BOS1), MYB112 and MYB116), their predicted lettuce orthologues (Reyes Chin-Wo et al, 2017), a cotton orthologue GhMYB108 (Genbank: ALL53614.1) and MYB124 (FLP) as an outgroup . **(B)** *B. cinerea* (top panels) and *S. sclerotiorum* (bottom panels) time series log2 expression (TPM) profiles of lettuce BOS1 orthologues (*Lsat_1_v5_gn_6_70301* and *Lsat_1_v5_gn_117600*), with pathogen inoculated expression profiles shown in red, and mock inoculated expression shown in blue. Shaded region shows 95% confidence intervals. **(C)** Expression of the *LsBOS1* gene (*Lsat_1_v5_gn_6_70301*) normalised to the *AtPUX1* housekeeping gene in Col-0 wildtype Arabidopsis and two independent lines of Col-0 expressing *LsBOS1* under control of the 35s promoter. Tissue was collected from pooled samples of 10 day old Arabidopsis seedlings with 3 technical replicates.

**Supp Figure 9:**
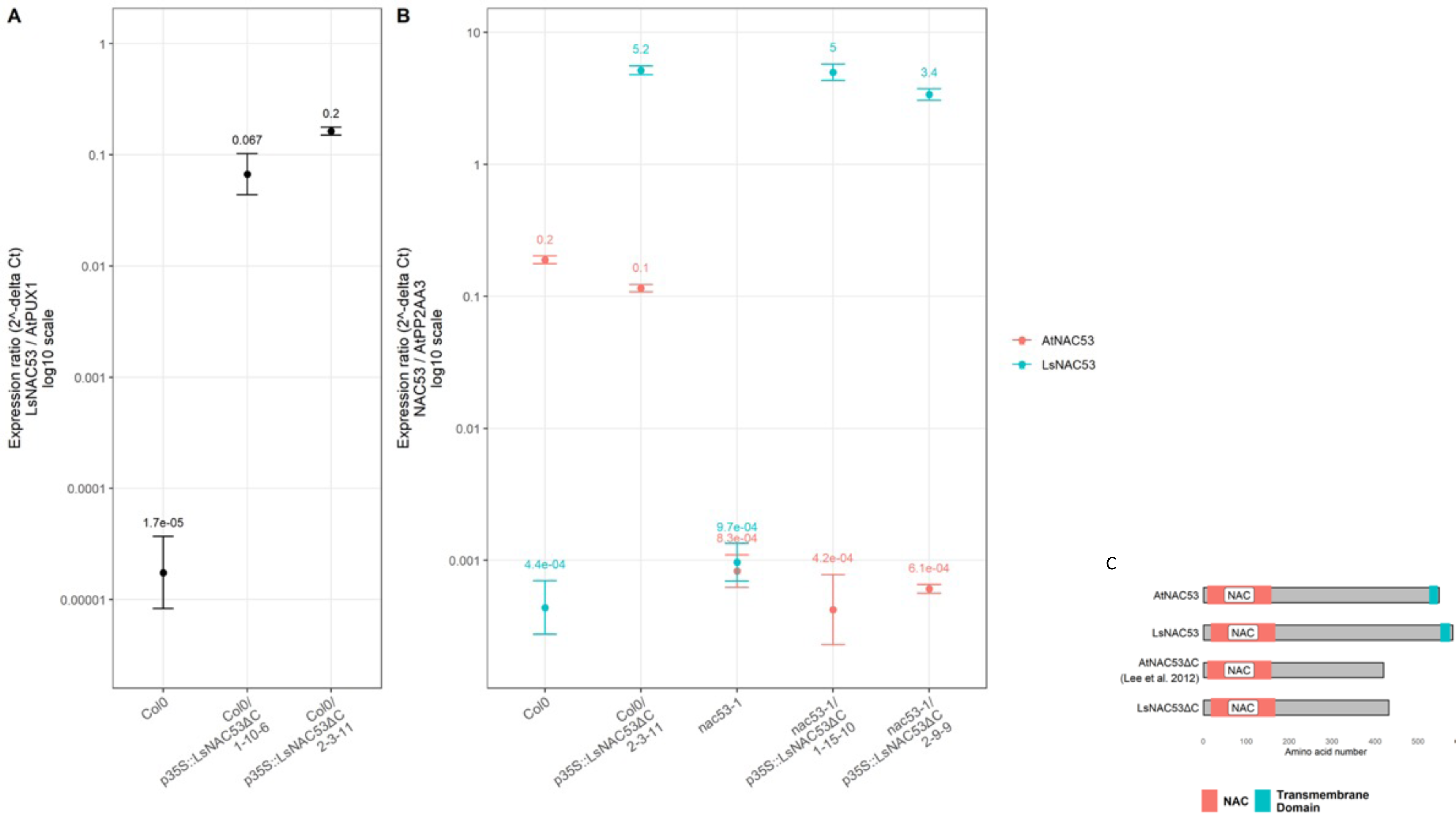
Characterisation of Arabidopsis transgenic lines expressing lettuce NAC53 orthologue. A) Expression of the *LsNAC53* gene (normalised to the *AtPUX1* housekeeping gene) in Col-0 wildtype Arabidopsis and two independent lines of Col-0 expressing *LsNAC53ΔC* under control of the 35s promoter. Both lines show a similar level of *LsNAC53* expression. B) Expression of the *LsNAC53* gene and endogenous *AtNAC53* (normalised to the *AtPP2AA3* housekeeping gene) in Col-0 wildtype Arabidopsis, one line of Col-0 expressing *LsNAC53ΔC* under control of the 35s promoter, the *nac53-1* mutant and two independent lines expressing *p35s::LsNAC53ΔC* in the mutant background. Endogenous *AtNAC53* expression is as expected in the wildtype and mutant lines and does not change in the presence of *LsNAC53*. All transgenic lines expressing *LsNC53* have a similar level of expression. Samples are pooled 10-day old Arabidopsis seedlings. C) The domain structure of NAC53, showing the N-terminal NAC DNA-binding domain and the C-terminal transmembrane domain, which prevents nuclear localisation. The truncation of LsNAC53*ΔC* is shown compared to the Arabidopsis truncated version used in Lee et al. 2012.

**Supplemental Figure 10:**
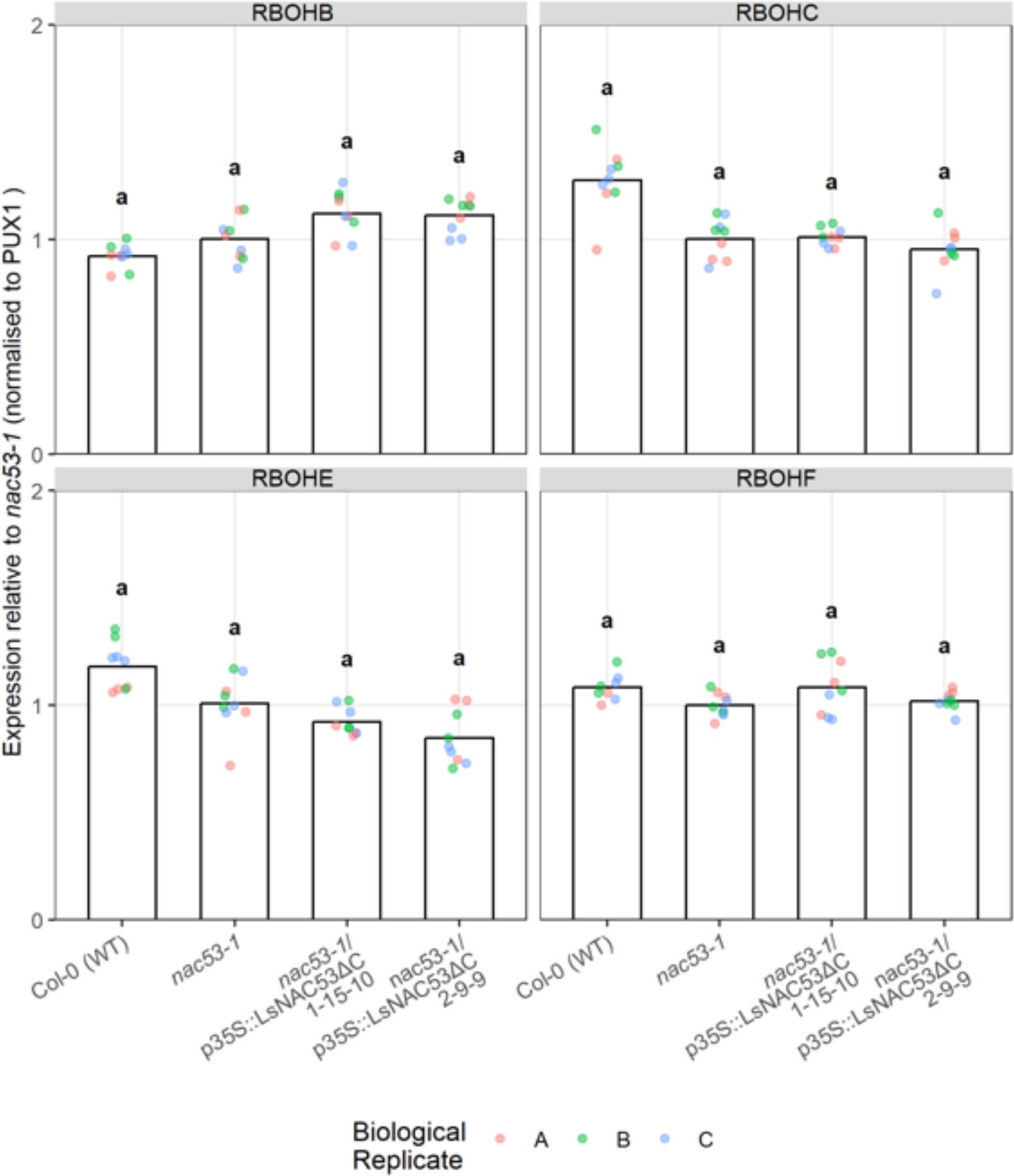
Relative expression of Arabidopsis genes RBOHB, RBOHC, RBOHE, RBOHF L1 in wildtype Col-0, *nac53-1* mutant and transgenic Arabidopsis *nac53-1* mutants expressing truncated *LsNAC53* under control of the 35s promoter (*nac53-1/ p35S::LsNAC53ΔC).* Three technical replicates of three biological replicates are shown with expression normalised to that of AtPUX1 and shown relative to expression in *nac53-1*. No statistical difference was observed based on the threshold of Tukey’s HSD p<0.05 and fold-change >1.5 or fold change <0.667 (equivalent downregulated fc).

